# Longitudinal clonal tracking in humanized mice reveals sustained polyclonal repopulation of gene-modified human-HSPC despite vector integration bias

**DOI:** 10.1101/2020.08.21.261537

**Authors:** Gajendra W. Suryawanshi, Hubert Arokium, Sanggu Kim, Wannisa Khamaikawin, Samantha Lin, Saki Shimizu, Koollawat Chupradit, YooJin Lee, Yiming Xie, Xin Guan, Vasantika Suryawanshi, Angela P. Presson, Dong-Sung An, Irvin S. Y. Chen

## Abstract

**Background:** Current understanding of hematopoiesis is largely derived from mouse models that are physiologically distant from humans. Humanized mice provide the most physiologically relevant small animal model to study human diseases, most notably preclinical gene-therapy studies. However, the clonal repopulation dynamics of human hematopoietic stem and progenitor cells (HSPC) in these animal models is only partially understood. Using a new clonal tracking methodology designed for small sample volumes, we aim to reveal the underlying clonal dynamics of human cell repopulation in a mouse environment.

**Methods:** Humanized BLT (bone marrow-liver-thymus) mice were generated by transplanting lentiviral vector transduced human fetal liver HSPC (FL-HSPC) in NOD.Cg-*Prkdc*^*scid*^*Il2rg*^*tm1Wjl*^/SzJ (NSG) mice implanted with a piece of human fetal thymus. We developed a methodology to track vector integration sites (VIS) in a mere 25µl of mouse blood for longitudinal and quantitative clonal analysis of human HSPC repopulation in mouse environment. We explored transcriptional and epigenetic features of human HSPC for possible VIS bias.

**Results:** 897 HSPC clones were longitudinally tracked in BLT mice—providing a first-ever demonstration of clonal dynamics and competitive expansion of therapeutic and control vector-modified human cell populations simultaneously repopulating in the same humanized mice. The polyclonal repopulation stabilized at 19 weeks post-transplant and the contribution of the largest clone doubled within 4 weeks. Moreover, 550 (∼60%) clones persisted over 6 weeks and were highly shared between different organs. The normal clonal profiles confirmed the safety of our gene therapy vectors. Multi-omics analysis of human FL-HSPC revealed that 54% of vector integrations in repopulating clones occurred within ±1kb of H3K36me3-enriched regions.

**Conclusions:** Human repopulation in mice is polyclonal and stabilizes more rapidly than that previously observed in humans. VIS preference for H3K36me3 has no apparent negative effects on HSPC repopulation. Our study provides a methodology to longitudinally track clonal repopulation in small animal models extensively used for stem cell and gene-therapy research and with lentiviral vectors designed for clinical applications. Results of this study provide a framework for understanding the clonal behavior of human HPSC repopulating in a mouse environment, critical for translating results from humanized mice models to the human settings.

## Background

Hematopoietic stem cells (HSC) are an ideal vehicle for introducing gene-modified cells to treat genetic disorders, cancers, and viral infections. Humanized mouse models—immunodeficient mice transplanted with human stem cells or tissues that generate a functioning human immune system—provide the most practical in vivo system for human stem cell and disease research (reviewed in [1, 2]). In particular, humanized bone marrow-liver-thymus mouse (hu-BLT mouse) models can support the development of human T cells, B cells, monocytes, macrophages, and dendritic cells. Comprehensive analysis of human T-cell development, diversity, and function in hu-BLT mice found intact TCRβ sequence diversity, thymic development, and differentiation to memory and effector cells [3]. The hu-BLT mice demonstrate human MHC-restricted T cell response to Epstein-Barr virus (EBV) infection and human dendritic cells-mediated T cell response against toxic shock syndrome toxin 1 (TSST1) [4]. The hu-BLT mouse models have been used in the field of cancer research to explore treatment strategy for non-Hodgkin’s lymphoma [5]. As models for sepsis studies, hu-BLT mice had high levels of human IL-6 in sera after cecal ligation and puncture (CLP)-surgery and are more susceptible to CLP induced sepsis than C57BL/6 mice[6]. A triple knockout (C57BL/6 Rag2^-/-^γc^-/-^CD47^-/-^) -bone marrow, liver, thymus (TKO-BLT) humanized mouse model was able to recapitulate the early in vivo human immune response against Ebola virus (EBOV) and Marburg virus (MARV) [7]. The Human Cytomegalovirus (HCMV) latently-infected hu-BLT mice developed human effector and central memory CD4+ and CD8+ T-cell responses and HCMV specific IgM and IgG B-cell responses [8]. Thus hu-BLT mice provide a valuable animal model system to study virus induced cellular and humoral immune response and to evaluate anti-viral treatments. To mimic anti-HIV gene-therapy in HIV patients, a recent study used an HIV-1 pre-infected hu-BLT mouse model and demonstrated that HIV-1 infection induces selective expansion of anti-HIV-1 dual shRNA gene-modified (protected) CD4+ T cells over control vector-modified unprotected CD4+ T cells [9]. Another recent study utilized HIV-infected hu-BLT mouse models to iteratively test CD28 and 4-1BB costimulation for optimizing anti-HIV CAR T cell therapy [10]. These studies reaffirm that hu-BLT mouse model provides an invaluable in vivo system for cancer, virus, and stem cell research and is particularly indispensable for preclinical gene therapy studies. However, translating results from xenograft mouse models to human setting is challenging as it remains unclear whether the human HSPC in mouse environment exhibit their human traits or clonally behave like mouse HSPC.

Understanding the clonal behavior of human cells in mouse environment is essential for interpreting results of humanized mice studies and effect of gene modification on the HSPC repopulation. In humans and macaques, longitudinal clonal tracking of transplanted HSPC revealed a biphasic expansion: an early phase of rapid and transient expansion of short-term HSC and a late phase (∼1 year post-transplant) of sustained expansion of long-term HSC [11, 12]. In mice transplanted with limited number of vector transduced mouse HSC, clonal tracking showed that clones start to stabilize around week 12 post-transplant and progressively fewer clones contribute to the overall repopulation [13]. Clonal tracking in mouse bone marrow also indicated similar time scale for clonal stabilization [14]. Another clonal tracking study in mice suggested transplantation dose-dependent change in HSC differentiation [15]. Clonal tracking in these mouse studies is done using genetic barcodes introduced in HSC using lentiviral vectors. Mice were transplanted with a limited number of mouse HSC that were transduced with lentiviral vector library consisting of individual vectors having a unique sequence tag or barcode. Integration of a barcoded vector in the DNA of host HSC introduces a unique genetic tag that can be used to identify and track individual HSC clones. This genetic barcoding assays enable quantitative clonal analysis and have been useful to study hematopoiesis and post-transplant clonal repopulation. However, generating a well curated barcode library for quantitative clonal tracking is both expensive and time consuming. Moreover, low DNA availability, lack of a universal barcode counting method, small barcode library size and other limitations [16] affect the accuracy and usability of vector barcoding techniques. Inability to identify genomic location of inserted barcode limits the use of genetic barcoding techniques to detect insertional mutagenesis induced clonal expansion.

Vector integration site (VIS) assays take advantage of a unique vector-host DNA junction sequence in each transduced HSPC due to the vector randomly integrating into host genome of these cells. A high-throughput integration sites (IS) sequencing assay can simultaneously identify multiple VIS and can quantitate relative contribution of each VIS clone to detect probable vector insertion induced clonal expansion. Our quantitative high-throughput VIS tracking study revealed that of all HSPC transplanted in rhesus macaques, ∼0.01% are long-term HSC and start contributing >1.5 years post-transplant [12]. Recently, our VIS assay found polyclonal repopulation of anti-HIV CAR modified HSPC in peripheral blood of pigtail macaques [17] indicating normal clonal expansion of stem cell derived CAR cells. Our long terminal repeat indexing–mediated integration site sequencing (LTRi-seq) method now enables multiplexed and unbiased quantitative clonal analysis of cells gene-modified with anti-HIV or control vector and that of HIV-1 IS—all in the same hu-BLT mouse [18]. Clonal analysis of different tissues/organs showed HIV-1 infection induced selective clonal expansion in the anti-HIV (H1 LTR-index) gene-modified population without adverse impact on clonal expansion of the control (H5 LTR-index) vector-modified population [18]. While these results show the potential of an anti-HIV gene-therapy, the underlying dynamics of clonal selection remain unknown owing to lack of a reliable quantitative clonal tracking technique for small animal models.

Although self-inactivating lentiviral vectors are low risk, a strong promoter within the vector can upregulate the expression of endogenous genes where the vector integrated [19, 20]. In rhesus macaques use of lentiviral vector containing a strong murine stem cell virus (MSCV) constitutive promoter-enhancer in the LTR induced aberrant clonal expansion [21]. Thus, promotor and transgene in vector can adversely impact the clonal behavior of transduced cells despite no obvious genotoxicity. Longitudinal clonal tracking of gene-modified cells in an appropriate small animal model. An in vitro study found lentiviral vector integration preference for active genes [22] and specifically associated with H3K36me3 [23] in activated human CD34+ HSC. In absence of appropriate in vivo longitudinal clonal tracking and vector integration data, the association between VIS preference for genes and histone modifications and its impact on in vivo proliferation, differentiation, and repopulation of gene-modified human HSPC remains partially explored.

Developing a reliable quantitative VIS assay for longitudinal clonal tracking in small animal models is challenging due to low sample volume. Only 100µl blood/biweekly can be safely drawn from a typical humanized mouse (∼ 0.6µg of DNA assuming 1000 cell/µl), whereas VIS assays require more than 1µg DNA. Multiple displacement amplification (MDA), a whole genome amplification technique, can increase the DNA amount from few nano grams to micro grams with a very low error rate (1 in 10^6^ to 10^7^ nucleotides) [24] and high coverage [25]. MDA amplified DNA has been used for various sequence sensitive applications such as single nucleotide polymorphism (SNP), next generation sequencing studies [26], detection of retroviral IS, and sequencing full-length HIV-1 proviruses including the IS [27, 28]. We hypothesize that MDA amplified DNA from low volume samples can be used with VIS assay for quantitative longitudinal clonal tracking in hu-BLT mouse. Thereby, providing an essential tool to study the clonal dynamics of HSPC repopulating in a mouse environment, to assess the safety of gene-therapy vectors, and to explore the association between in vivo clonal behavior of gene-modified HSPC and location of vector integration in host genome.

In this study, we developed a quantitative low volume vector integration site sequencing (LoVIS-Seq) assay, a combined MDA and VIS assay for low-volume samples, for longitudinal clonal tracking in small animal models. We longitudinally tracked hundreds of clones in two different gene-modified cell populations simultaneously repopulating in hu-BLT mice. Human cell repopulation in a mouse environment is polyclonal and resembles typical after-transplant HSPC expansion in other small and large animal models. Analysis of multi-omics data of FL-HSPC and VIS in repopulating clones shows vector integration bias for actively transcribed regions of the genome. Our assay provides an efficient tool to study clonal repopulation in murine and humanized-mouse models used for stem cell and gene therapy research. Results of clonal and genomic analysis revealed valuable insight into the clonal dynamics of human HSPC repopulation in a mouse environment.

## Methods

### Human fetal thymus and isolation of FL-CD34+ cells from fetal tissue

Human fetal thymus and livers were obtained from Advanced Bioscience Resources (ABR) and the UCLA CFAR Gene and Cellular Therapy Core. Human fetal liver CD34+ HSPC and thymus pieces were processed as previously described [29]. Briefly, a single cell suspension of fetal liver cells was strained through 70µm mesh and layered onto density gradient separation media (Ficol Paque PLUS, GE Healthcare). After 20 minutes of centrifugation, the mononuclear cells layer was collected. Anti-CD34+ microbeads (Miltenyi Biotech, San Diego, CA) were used for magnetic isolation of CD34+ cells from mononuclear cells. Calvanese et al. [30] also obtained fetal liver from the UCLA CFAR Gene and Cellular Therapy Core and followed identical CD34+ magnetic sorting to isolate uncultured FL-HSPC for RNA-seq, ATAC-seq, and ChIP-seq assay.

### Vector transduction of FL-CD34+ cells

The isolated FL-CD34+ cells were transduced overnight using either mCherry or EGFP vector at MOI (multiplicity of infection) of 1 and 3, respectively. We used higher MOI for EGFP vector to achieve gene marking level comparable to mCherry vector. Fraction of vector transduced cells were cultured for 4 days and percentage of EGFP or mCherry expressing cells were measured by flow cytometry (Supplementary figure 2A). Our data showed of the FL-CD34 cells transduced with mCherry vector, 68.4% cells were mCherry+. For EGFP vector of the FL-CD34 cells transduced with EGFP vector 88.5% cells were EGFP+.

### Humanized BLT mouse and sample collection

NOD.Cg-*Prkdc*^*scid*^*Il2rg*^*tm1Wjl*^/SzJ (NSG) mice, 6-8-week-old, were myeloablated 1 day before transplant by intraperitoneal (i.p.) injection with 10 mg/kg of 6-thioguanine (6TG) (Sigma-Aldrich, Saint Louis, MO) or 35mg/kg of Busulfan for mouse m860. Myelo-preconditioned mice were transplanted with human fetal liver CD34+ HSPCs transduced with Anti-HIV vector (EGFP+) (0.5 × 10^6^ cells/mouse) and mixed with HSPCs transduced with the control (mCherry+) vector (0.5 × 10^6^ cells/mouse). The mice were transplanted with a two-step procedure: half the mixture of EGFP+ and mCherry+ transduced cells was solidified by matrigel (BD Bioscience, San Jose, CA), mixed with CD34-cells as feeder cells (4.5×10^6^ cells), and implanted with a piece of human thymus under the mouse kidney capsule. Then, mice were injected with the other half of the mixed EGFP+ and mCherry+ transduced cells via retro-orbital vein plexus using a 27-gauge needle on the same day. Bone marrow cells for MDA were harvested from mouse m860 at week 25 post-transplant. Low human reconstitutions at early timepoints and graft versus host-disease related illnesses at very late timepoints post-transplant have potential to impact interpretation of the clonal data. In order to avoid these potential confounding effects, we longitudinally tracked clones between week 13 to 19 post-transplant. Every two weeks from weeks 13-19 post-transplant, 100µl of mouse blood was collected from the retro-orbital vein for longitudinal clonal tracking and for monitoring reconstitution of human leukocyte and level of the EGFP+ and mCherry+ marked cells. Plasma was removed and peripheral blood cells were stained with monoclonal antibodies for 30 minutes (Supplementary figure 9). Red blood cells were lysed with red blood cell lysis buffer (4.15 g of NH4Cl, 0.5 g of KHCO3, and 0.019 g of ethylenediaminetetraacetic acid in 500 mL of H2O) for 10 minutes and washed with FACS buffer (2% fetal calf serum in phosphate-buffered saline [PBS]). Stained cells were resuspended in 16µl PBS, of which 8µl was split equally into two tubes for MDA replicates. The remaining 8µl was mixed with 300µl of 1% formaldehyde in PBS and examined with Fortessa (BD Biosciences) flow cytometers. Flow cytometry data was utilized to monitor human reconstitution (Supplementary table 2) and count human cells, mCherry+ cells, and EGFP+ cells as well as human T and B cells in blood (Supplementary figure 2B). The following monoclonal antibodies with fluorochromes were used: human CD45-eFluor 450 (HI30, eBioscience), CD3-APC-H7 (SK7: BD Pharmingen), and CD19-BV605 (HIB19: BioLegend). Data were analyzed on FlowJo (TreeStar, Ashland, OR) software.

The UCLA Institutional Review Board has determined that fetal tissues from diseased fetuses obtained without patient identification information are not human subjects. Written informed consent was obtained from patients for use of these tissues for research purposes. All mice were maintained at the UCLA Center for AIDS Research (CFAR) Humanized Mouse Core Laboratory in accordance with a protocol approved by the UCLA Animal Research Committee.

### LoVIS-Seq workflow with whole genome amplification and quantitative VIS assay

Multiple displacement amplification for whole genome amplification: To estimate the minimum number of cells required for LoVIS-Seq, we collected 81,000, 27,000, 18,000, 9,000, 3,000, and 1,000 bone marrow cells from mouse m860 by serial dilution and stored in 4µl of PBS at -20ºC. MDA was done directly on cells (Supplementary figure 9) using the REPLI-g Single Cell Kit from Qiagen (Cat #150343) following kit-specific protocol. For longitudinal clonal tracking in the blood compartment, 100 µl blood was drawn at weeks 13, 15 & 17. At end point (week 19), max blood (≈ 1ml) was collected, out of which 100µl was used for flow cytometric analysis along with MDA; the remainder was used to isolate unamplified whole blood DNA. Cells for MDA were isolated as described above and stored in 4µl of PBS at -20ºC. MDA-amplified DNA was then used for quantitative VIS assay. A Qiagen DNeasy Blood & Tissue Kit was used to extract unamplified DNA from max blood cells, splenocytes, and bone marrow cells.

Quantitative VIS assay and data analysis workflow: For VIS sequencing, we followed the procedures described in our previous publication [12, 18, 31, 32] and focused on analyzing only the right LTR junctions using CviQI and RsaI restriction enzymes. For our VIS assay, we used one microgram MDA-amplified or unamplified genomic DNA for animal m860 samples and two micrograms MDA-amplified or unamplified genomic DNA for different time point samples, with a few exceptions (see Supplementary Tables 1 and 2). DNA samples were subject to extension PCR using LTR specific biotinylated primers /5BiotinTEG/CTGGCTAACTAGGGAACCCACT 3’ and /5BiotinTEG/CAGATCTGAGCCTGGGAGCTC 3’. The extension PCR product was then digested using CviQI and RsaI restriction enzymes and biotin primer bound DNAs isolated using streptavidin-agarose Dynabeads using magnetic separator as per manufactures instructions. The vector-host junctions capture on streptavidin beads were processed for linker-mediated PCR (LM-PCR) methods as described previously[31, 32]. The linker ligated vector-host junction DNA was subjected to two step PCR. First step amplification was done using primer 5’ CTGGCTAACTAGGGAACCCACT 3’ and first linker primer GTGTCACACCTGGAGATAT. We removed the internal vector sequence by restriction enzyme (SfoI) digestion. The digested product of first PCR was then amplified using primer 5’ACTCTGGTAACTAGAGATCC 3’ and second linker primer 5’ GGAGATATGATGCGGGATC 3’. Since the LTR index sequence is included in the vector-host junction, we obtain unbiased amplification all the H1, H5 and/or WT VIS sequences. Lentiviral vectors used in this study as derived from FG12-mCherry lentiviral vector[29] and all the primers are designed accordingly. A detailed protocol for VIS assay is provided in supplementary text. The amplicon libraries prepared using custom made Illumina (Illumina, San Diego, CA) sequencing primers for Illumina MiSeq (m860 samples) or iSeq100 (m599, m599, and m591 samples) sequencer. Sequences with a virus-host junction with the 3’ end LTR, including both the 3’-end U5 LTR DNA and ≥ 25 base host DNA (with ≥ 95% homology to the human genome), were considered true VIS read-outs. The sequence mapping and counting method was performed as described previously[18]. In brief, sequences that matched the 3’end LTR sequence joined to genomic DNA as well as LTR-indexes (H1-TGGAAAATCTCCAACA, H5-TGGAAAATATCCAACA or WT-TGGAAAATCTCTAGCA) were identified using a modified version of SSW library in C++ [33]. Reads were classified as H1, H5, or WT VIS based on the LTR barcodes used in the experiment. VIS sequences were mapped onto the human genome (Version hg38 downloaded from https://genome.ucsc.edu/) using Burrows-Wheeler Aligner (BWA) software. Mapped genomic regions were then used as reference and VIS reads were remapped using BLAST to further remove poorly mapped reads to get an accurate estimate of sequence count. Final VIS counting was done after correcting for VIS collision events and signal crossover as described previously. VIS with a final sequence count less than the total number of samples analyzed per animal were removed. VIS clones with maximum frequency values below 1^st^ quartile were classified as “low frequency”, clones with maximum frequency value above 3^rd^ quartile were classified as “high frequency”, and clones with maximum frequency between the 1^st^ and 3^rd^ quartiles were designated “medium frequency”. VIS clones that were detected with frequency >0 at every week from 13-19 are termed “persistent clones”. The 10 high frequency VIS clones at each timepoint were selected as top 10 VIS. All the VIS data and list of VIS-proximal genes is provided in supplementary file.

### Random integration sites

Random integration sites were generated in silico using a custom python script. To mimic our VIS assay, we randomly selected 1000 integration sites that were within ±1500bp of the nearest CviQI/RsaI (GTAC) site in the human genome (hg38).

### Clonal diversity analysis

For diversity analysis, we used Rényi’s diversity/entropy [34] of order *α* defined as follows

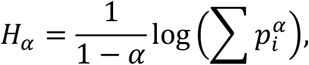

Where *p*_*i*_ is the proportional abundance of the *i*th VIS clone for *i = 1, …, n*. At each timepoint, an average Rényi’s diversity profile was obtained by calculating average values of ***H***_***α***_ for *α* ≥ 0. The *α* is considered as a weighting parameter such that increasing *α* leads to increased influence of high frequency VIS clones. The proportional abundance is calculated as *p*_*i*_ = *s*_*i*_*/S*, where *s*_*i*_ is the sequence count of the *i*th VIS clone and *S* is the sum of sequence counts from all VIS clones. The Rényi’s diversity ***H***_***α***_ values are averaged over two replicates and plotted as a function of *α*. If all VIS clones contributed equally, i.e. 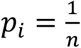 for all *i* = 1, …, *n*, then ***H***_***α***_ for all values of *α* would be equal and the profile (line) would be horizontal. VIS clones expanding at different rates would show decreasing ***H***_***α***_ values as *α* increases, generating a downward-sloped diversity profile that is steeper with more non-uniform clonal expansion. ***H***_***α***_ indicates clonal diversity of the repopulating cells, such that consistently higher values of ***H***_***α***_ indicate a more diverse clonal population. If the profiles for two populations/samples cross, then their relative diversities are similar. For *α* = 0, ***H*_*0*_** = *log(n)* and the antilogarithm of this value equates to the richness or number of unique IS. ***H***_***α***_ at *α* = 1 and *α* = 2 are the Shannon and 1/Simpson indexes, respectively. We calculated Renyi’s diversity using the R package BiodiversityR (*https://cran.r-project.org/web/packages/BiodiversityR/index.html*). For the above analysis, we used raw sequence counts from two replicates without distinguishing between mCherry-H5 VIS and EGFP-WT VIS.

### RNA-seq data analysis

Raw sequence data of uncultured FL-HSPC (in triplicate) was pre-processed for quality using Fastqc. Trimmomatic was used to remove adaptors and for quality trimming. After this, reads were aligned onto human genome hg38 using RNA STAR aligner [35]. SAMtools was used to remove reads with low mapping scores (< 20) and to generate BAM files. Cufflinks [36] was used to calculate FPKM values for all genes. The human cancer consensus gene list is from Catalogue of Somatic Mutations In Cancer (https://cancer.sanger.ac.uk/census).

### ATAC-seq analysis

Raw sequence data of uncultured FL-HSPC (in triplicate) was pre-processed for quality using Fastqc. Adaptor removal and quality trimming was done using Trimmomatic. After this, reads were mapped onto human genome hg38 using bowtie2 with parameter --very-sensitive -X 2000 -k 1. SAMtools was used to remove reads with low mapping (<20) scores, blacklisted regions, and to generate BAM files. Picard tool kit was used to remove duplicate reads. We used Genrich, a paired end peak caller, to identify ATAC peaks. Software deepTools[37] was used to generate coverage (.bw) files and for visualization of open DNA in genes and VIS-proximal regions.

### ChIP-seq data analysis

Raw sequence data of uncultured FL-HSPC for histones, RNAploII, and input were pre-processed for quality using Fastqc. Trimmomatic was used to remove adaptors and for quality trimming. After this, reads were mapped onto human genome hg38 using bowtie2 with parameter --local. SAMtools was used to remove reads with low mapping (<20) scores, blacklisted regions, and to convert SAM to BAM format. Picard tool kit was used to remove duplicates. MACS2 tool was used to call peaks for all histone marks and RNApolII using input sample as control. Software deepTools [37] was used to generate coverage .bw files and for visualization of histone/RNApolII in genes and VIS proximal regions. Proximity of VIS to a peak was calculated by measuring the distance between VIS and the coordinates of the nearest peak; for VIS within the peak distance is 0bp.

### Statistical analysis

Relative clonal frequencies are summarized as means ± standard deviations (SDs). Pearson correlation (r) is used to assess similarity in terms of a linear relationship among pairs of samples. The interclass correlation coefficient, ICC(3,1) two-way mixed effects, absolute agreement is used to compare the reproducibility (ICC) of clonal profiles among replicates [38]. ICC, Pearson’s r and p values are calculated using statistical software R (version 3.6, https://www.r-project.org/). To determine if VIS preference for genomic and epigenetic features differs significantly from random IS, we used Pearson’s chi-squared test with Yate’s continuity correction (function chisq.test() in software R). We used Principal Component Analysis (PCA) to reduce the complexity of read coverage data of multiple chromatin feature in proximity to VIS. The dimensionality reduction by PCA method is similar to clustering and allows detection of patterns in the data. In this study, PCA was done using software deepTools [37].

## Results

### Minimum 10,000 bone marrow cells or 25µl blood is sufficient for LoVIS-Seq

To estimate the minimum number of cells sufficient for LoVIS-seq assay to provide a quantitative clonal analysis, we collected bone marrow (BM) cells from hu-BLT (bone marrow-liver-thymus) mouse (m860). Fetal liver CD34+ cells transduced with sh1005(anti-CCR5 shRNA)-EGFP vector or control mCherry vector were mixed in equal ratio and transplanted in the mouse (Figure 1a-b, details in Methods). We first estimated clonal composition of the EGFP-WT (WT LTR-index) and mCherry-H1 (H1 LTR-index) populations using unamplified bulk DNA, in triplicate, as described previously [18]. A total of 300 ±42 SD (216 ±22 SD mCherry-H1 VIS and 84 ±20 SD EGFP-WT) VIS were detected indicating polyclonal expansion. The clonal size distribution in the mouse bone marrow (Figure 1c) resembled that of recently observed hu-BLT mice [18] and pigtail macaques [17] by us as well as that of observed by others in mice [13], autologously transplanted nonhuman primates [12, 39], and humans [11, 40] overall, indicating normal clonal repopulations.

**Figure 1:**
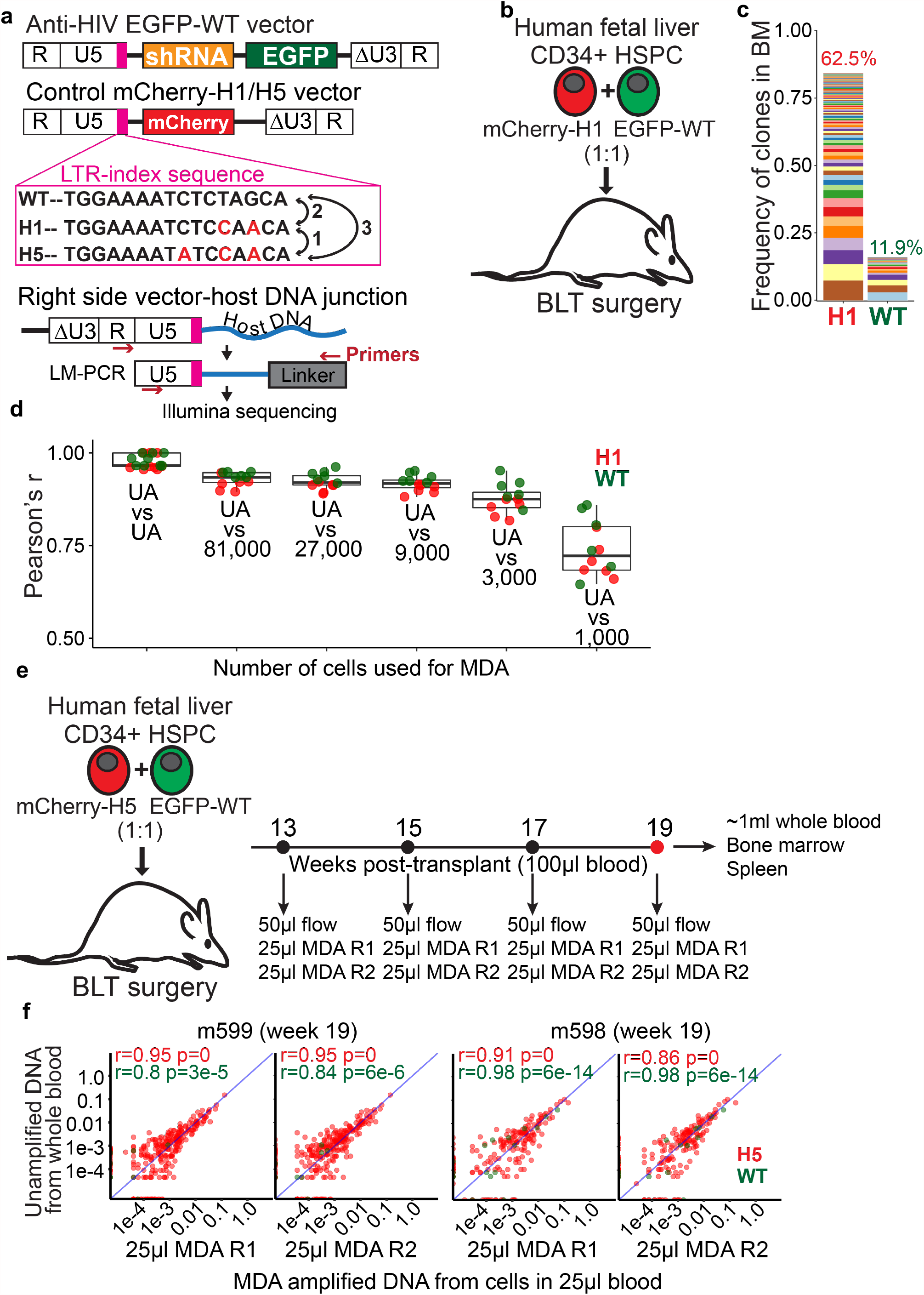
LoVIS-Seq reproduces clonal distribution of entire mouse blood using 25µl blood: **a**) Diagram showing Anti-HIV-EGFP-WT and control mCherry-H1/H5 vectors having WT, H1, or H5 LTR-index and strategy for VIS assay with LTRi-seq. LTR index sequences are shown in magenta box. **b**) Hu-BLT mouse model: Fetal liver CD34+ cells were separately transduced with either anti-HIV or control vectors and transduced cells were mixed 1:1 for transplant. The mix of transduced cells was transplanted in myeloablated NSG mice with a fetal thymus tissue implant. **c**) Stacked bar plot showing clonal frequencies of VIS in BM of hu-BLT mouse. Clones from mCherry-H1 and EGFP-WT cells were identified by corresponding LTR barcodes. In the stacked bar plot, each band represents a unique VIS (HSPC clone) and thickness of the band shows clonal frequency or abundance of that HSPC clone. Percentage of mCherry+ or EGFP+ cells within human cell (hCD45+) population are shown on top of the corresponding stacked-bar. **d**) Plot showing Pearson’s r for correlations of mCherry-H1 (red dots) and EGFP-WT (green dots) VIS clonal profiles between unamplified DNA replicates and replicates of MDA-amplified DNA samples for different cell numbers. **e**) Experimental protocol for longitudinal clonal tracking in humanized BLT mice. **f**) Scatter plot showing VIS clonal frequencies between unamplified whole blood DNA and two replicates of MDA-amplified DNA from 25µl blood at week 19 (r= Pearson’s r, diagonal line is r=1) for m599 and m598. Clonal frequency of mCherry-H5 (red dots) and EGFP-WT (green dots) VIS clones in unamplified DNA samples (y-axis) and MDA replicates (x-axis).

For low volume sample with DNA yield of <1µg, we propose MDA to increasing the quantity of and use the amplified DNA for VIS assays. For our LoVIS-seq assay, we first perform MDA directly on the cells and use the MDA amplified DNA for VIS assays (Supplementary figure 9). We hypothesized that LoVIS-seq can be used for clonal analysis of low volume samples without loss of quantitative accuracy. To test our hypothesis, we first performed MDA directly on 81,000, 27,000, 9,000, 3,000, and 1,000 bone marrow cells, each in duplicate (Supplementary figure 1A, details in Methods section). We used equal amounts of MDA-amplified DNA and unamplified bulk DNA for the VIS assay (Supplementary table 1). We assessed reproducibility of clonal profiles in MDA amplified DNA of 81,000, 27,000, 9,000, 3,000, and 1,000 bone marrow cells against that of unamplified DNA sample from bulk BM cells. We performed routinely used Pearson correlation analysis and also did Intraclass Correlation Coefficient (ICC) analysis to test the reproducibility of clonal abundances (details in Method section). We found high reproducibility of clonal profiles in different MDA samples and within-MDA replicates of 81,000 to 9,000 cells (avg. Pearson’s r value >0.91) (Figure 1d and Supplementary figure 1B-F). To determine the minimum number of BM cells that are required for obtaining high quantitative accuracy, we compared the reproducibility of MDA amplified 81,000, 27,000, 9,000, 3,000, and 1,000 bone marrow cells and found that for less than 9,000 cells, the reproducibility dropped (avg. Pearson’s r=0.87 for 3,000 and 0.73 for 1,000 cells). Importantly, reduced cell numbers caused a modest reduction in VIS detection (Supplementary figure 1G) showing minimal impact on the sensitivity of our assays. In order to further validate the reproducibility of our assay, we used ICC analysis. ICC can be used to simultaneously compare replicates of unamplified DNA against replicates of MDA amplified samples. The ICC values (Supplementary figure 1H) were similar to Pearson’s r, further indicating loss of reproducibility when fewer cells were used as source material for MDA. Overall, both the Pearson and ICC analysis, validate that MDA-amplified DNA from >10,000 bone marrow cells is sufficient for LoVIS-Seq.

After establishing the accuracy and reproducibility using BM cells, we tested LoVIS-seq assays with hu-BLT mouse blood. We collected 100µl blood at week 13, 15, 17 and ∼1ml of whole blood at week 19 post-transplant from hu-BLT mice. These mice were transplanted with an equal mix of human CD34+ cells transduced with anti-HIV EGFP-WT vector and control mCherry-H5 vector (Figure 1e). Cells from 50µl blood were used for flow cytometry and the remaining cells were used for MDA duplicates; each 25µl of blood (∼400 and ∼600 human cells/µl blood for mouse m599 and m598, respectively). High correlation (median Pearson’s r =0.93) of mCherry-H5 and EGFP-WT VIS clonal frequencies between unamplified and MDA-amplified DNA from blood cells (Figure 1f) indicates clonality of entire mouse blood can be captured with 25µl of blood. Importantly, the MDA replicates also showed high reproducibility (median Pearson’s r>0.95, Supplementary figure 3A). To further validate the reproducibility, we simultaneously compared two MDA replicates and unamplified DNA from whole blood. The ICC values of >0.95 further confirmed reproducibility of our assay with mere 25µl blood. In conclusion, our LoVIS-Seq assay accurately captured the clonality of two vector-modified cell populations in hu-BLT mouse blood using mere 25µl of blood or as few as 10,000 cells. These results indicate that LoVIS-Seq assay can be used to study clonal repopulation of gene-modified cells in mouse blood using only a fraction of (approximately 2.5%) of whole blood.

### Simultaneous clonal tracking of therapeutic vector-modified and control vector-modified populations

After demonstrating the high accuracy and reproducibility of LoVIS-Seq assay, we aim to examine clonal dynamics of anti-HIV EGFP-WT vector and control mCherry-H5 vector modified two human cell populations simultaneously repopulating the mouse blood. We collected 25µl of hu-BLT mouse blood samples for each MDA replicate on week 13, 15, 17 and 19 post-transplant. Despite the variation in cell count over time each replicate/sample had >10,000 (avg. 21,189; ±7,364) cells (Supplementary table 4 and Supplementary figure 2B). After performing LoVIS-Seq assay on these cell samples, we identified 329 VIS in m599, 222 VIS in m598, and 346 VIS in m591 (total 897 VIS clones) and longitudinally tracked change in the relative contribution of these clones to the repopulation from week 13 to 19 (Figure 2a). From total 897 VIS clones, we classified mCherry-H5 (792) and EGFP-WT (105) VIS clones using LTR-indexes in the two vectors. Consistent with our previous study [18], we observed high correlations between the total mCherry-H5 VIS clonal frequency and mCherry^+^ cell percentage by flow cytometry (Pearson’s r = 0.8) as well as between total EGFP-WT VIS clonal frequency and EGFP^+^ cells percentage (Pearson’s r = 0.9, Supplementary figure 3B). The high correlation between flow cytometry and clonal frequency data indicates quantitative accuracy of LoVIS-Seq assay and ability to simultaneously track clonal expansion of two human cell populations repopulating in the same animal using a mere 25µl of blood. In addition, longitudinal clonal tracking further shows expansion in EGFP-WT clones coincided with reduction of mCherry-H5 contribution and vice-versa (Figure 2b; solid lines); these changes closely match the change in EGFP^+^ and mCherry^+^ cell percentages measured by flow cytometry (Figure 2b; dashed lines). Furthermore, repopulation in both EGFP-WT and mCherry-H5 populations is largely driven by expansion of a few HSPC clones—a characteristic feature of after-transplant repopulation. These results present a first-ever demonstration of clonal expansion of two populations, therapeutic vector-modified and control vector-modified, competing to repopulate in hu-BLT mice.

**Figure 2:**
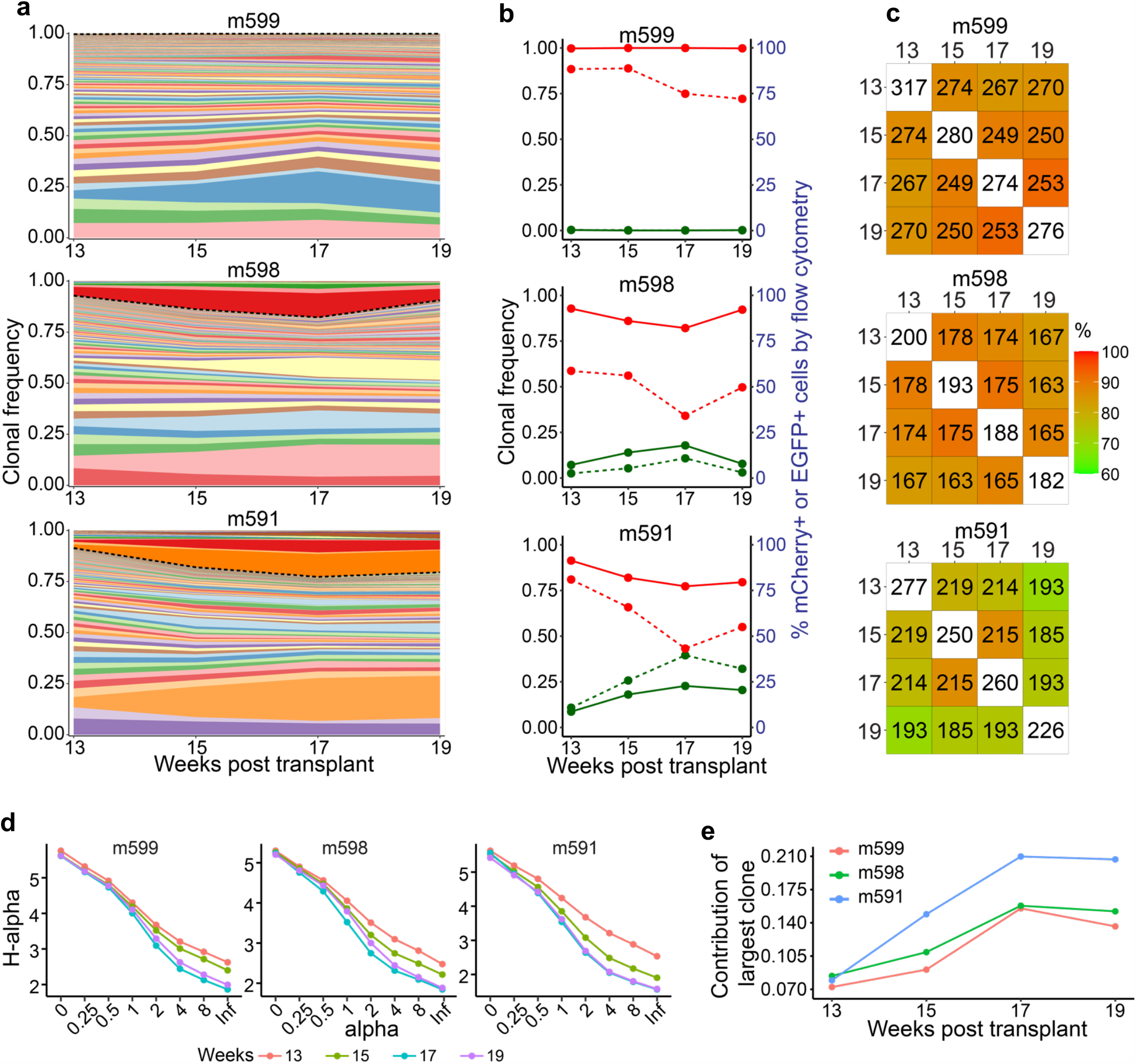
Longitudinal clonal tracking in hu-BLT mice: **a**) Area plots show clonal repopulation in whole blood over time from week 13-19. Each colored band is a unique VIS clone and thickness of the band corresponds to frequency of the VIS clone. Dashed black line separates mCherry-H5 VIS clones (below) and EGFP-WT VIS clones (above). **b**) Line plots show changes over time in the total frequency of mCherry-H1 VIS clones (solid red line) and total frequency of EGFP-WT VIS clones (solid green line) as well as percentages of mCherry+ cells (dashed red line) and EGFP+ cells (dashed green line). **c**) Heatmaps showing percentage change in shared clones between two timepoints. Digits inside white tiles on the diagonal show number of VIS detected at each time point. Colors of each heatmap tile correspond to percentage of clones shared and color key is provided on the right. Digits in each tile show the number of VIS shared between two timepoints. **d**) Renyi’s diversity profiles evaluated using raw count data from two replicates at each timepoint and by varying value of alpha. Renyi’s diversity profiles are arranged with highest diversity at the top to lowest at the bottom. Topmost curve with no overlap or intersection with any other curve has the highest overall diversity. Diversity of curves that overlap or intersect is undefined. **e**) Line plot showing contribution of highest contributing clone at different timepoints. Values reported are exp(H(alpha)) at alpha=∞.

### Sustained polyclonal repopulation with rapid clonal expansion and stabilization

The clonal analysis revealed that at each timepoint hundreds of VIS clones contributed to the repopulation and contributions of EGFP-WT and mCherry-H5 clones changed over time. To investigate the properties of the clonal dynamics of human cell repopulation in mouse environment, we analyzed the quantitative clonal tracking data from different timepoints. We found that the maximum number of VIS were detected at week 13 and on average, only 13% (±5%) fewer VIS were detected at week 19 compared to week 13. Also, the number of total VIS contributing to repopulation at each timepoint decreased with time (Figure 2c). On average, 61% (±12.8%) of persistent VIS clones (m599: 235 clones, m598: 150 clones, and m591: 165 clones; total 550 clones) consistently contributed for 6 weeks and provided stable polyclonal repopulation (Figure 2a area plots). While the number of VIS clones steadily dropped over time in both the mCherry-H5 and EGFP-WT populations, their clonal profiles became increasingly similar (Supplementary figure 4A-B), comparable to polyclonal repopulation patterns that have been reported in nonhuman primates [12]. Moreover, correlation between time points indicates clonal distribution at week 13 differs from week 19, with clonal expansion stabilizing around week 17 (Supplementary figure 4B). We also examined Rényi’s diversity profiles for each animal at each time point (details in Methods section). The clonal diversity at week 13 was highest and subsequently decreased with time (Figure 2d). Diversities were similar between weeks 17 and 19 as indicated by their overlapping diversity profiles. The two commonly used diversity indexes Shannon Index [41] (α = 1) and ln(1/Simpson index) (α = 2) referred here on as Simpson index [42] to capture the overall change in the diversity using single numerical values. Shannon index is more sensitive to number VIS clones and Simpson index to the relative frequency of each VIS clones. The Shannon[41] and Simpson[42] indices dropped between weeks 13 and 17, indicating expansion of fewer clones (Supplementary figure 4C). The similarity of the indices between weeks 17 and 19 suggests stabilization of clones. Contribution by the most frequent clone rose ∼2.2 times, from 0.078 (±0.005, n=3) at week 13 to 0.174 (±0.030, n=3) at week 17 (Figure 2e), signaling rapid expansion of a few clones. Compared to humans, wherein stable clones appear >1 year post-transplant [11], in hu-BLT mice the clonal repopulation remained normal despite faster expansion and earlier stabilization of human HSPC clones. Overall, these results indicate notable difference in the time scale of clonal repopulation of human cells in mouse from that in human environment and these finding have larger implication while interpreting and translating results from hu-BLT mouse models to human settings.

### Clonal sharing between organs reveals normal repopulation and unique tissue distribution pattern

We showed first-ever demonstration of clonal repopulation of two human cell population in mouse environment with sustained polyclonal repopulation and competitive expansion of two human cell population in mouse blood. However, clonal studies in nonhuman primates revealed that clonal expansion patterns in blood during early phase of repopulation differ from other organs [43]. In hu-BLT mice whether clonality of blood differs from the tissue/organ is unknown. We determined the clone profiles is bone marrow (BM) and spleens compartments of these mice using bulk cells for our VIS assay. Similar to blood, BM and spleen compartments also showed normal polyclonal repopulation. Between organ/tissue comparison of clonal size distributions showed a highly correlated clonal expansion pattern (avg. Pearson’s r =0.96) between blood and spleen (Figure 3); however, clonal expansion in bone marrow was less correlated to blood or spleen. Importantly, we observed that in all three tissue compartments, persistent clones contributed the most to repopulation. These results show substantial clonal sharing among yet, differential clonal contributions. Our data also indicate that clonal tracking using BM samples would reflect the clonality only of the BM compartment. Compared to serial bone marrow extractions, clonal tracking using blood is more practical and closely reflect clonal composition of spleen.

**Figure 3:**
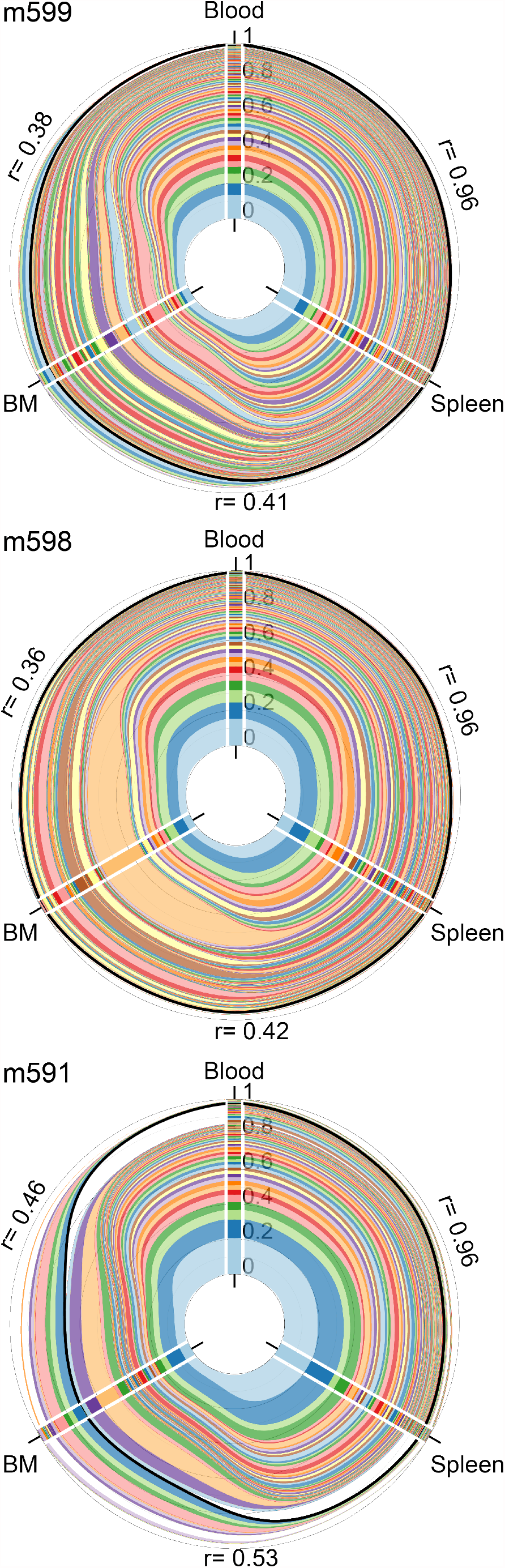
Unique clonal sharing pattern between different tissues: Polar area plots of clonal expansion and sharing in peripheral blood, spleen, and bone marrow (BM). There are three axes, one for each tissue. Stacked bar plot on each axis shows size distribution of clones in the tissue. Each colored stack represents a VIS clone and its thickness shows abundance of the clone. Clones shared between tissues are connected using ribbons with colors matching the clone’s stack color in the bar plot. Black line encompasses total size distribution of persistent clones that continuously contributed from week 13 till 19. Clones did not continuously contribute for 6 weeks and were detected at week 19 in blood, spleen, and BM are outside black line. Pearson’s r values are shown in black.

### Influence of genomic location and proximal genes on clonal growth

We found normal and polyclonal expansion in human cells in mouse blood and tissue compartment yet mutagenic insertions causing abnormal growth remains a major concern for gene-therapy vectors. For each VIS, our assay provided both relative frequency and genomic location of integration allowing us to identify possible of insertional mutagenesis induced clonal growth. Our VIS data from in vivo repopulating clones showed preference for high gene density chromosomes (Supplementary figure 5A) also observed with the HIV-1 integration pattern [44]. Additionally, similar genomic distribution of low, medium, and high frequency in vivo repopulating VIS clones (Figure 4a) suggests clonal expansion is unrelated to genomic location of integration. We found that in ∼80% of the total detected clones, VIS occurred within ±1Kb of protein-coding genes, significantly higher than 54% of 1000 random IS (p< 0.001, Supplementary figure 6A). About 8% of VIS were within ±1Kb of long non-coding RNA (lncRNA) and 10% were outside ±1Kb of any genes (distal VIS, Figure 4b inside pie chart). Persistent and top 10 VIS clones (top 10 high frequency VIS clones collected from each timepoint) showed similar preference for gene biotypes (Figure 4B inside pie chart). Out of all VIS, only 66 (including 1 out of 42 top 10 VIS) were proximal to known cancer consensus genes (Figure 4a Circos plot). Gene ontology analysis of proximal genes and their mouse orthologs showed significant enrichment (P < 0.01) in various biological pathways such as cell-cell interaction, viral process and transcription regulation (Supplementary figure 5B). In vitro data of VIS-proximal gene in vector transduced human CD34+ HSC showed enrichment in similar biological processes.[22] A recent study claims that biological function of genes near HIV integrations sites influence in vivo clonal expansion of HIV infected cells.[45] However, we found no strong evidence of link between in vivo differentiation potential of VIS clones and genomic location of its VIS or biological function of the VIS-proximal genes.

**Figure 4:**
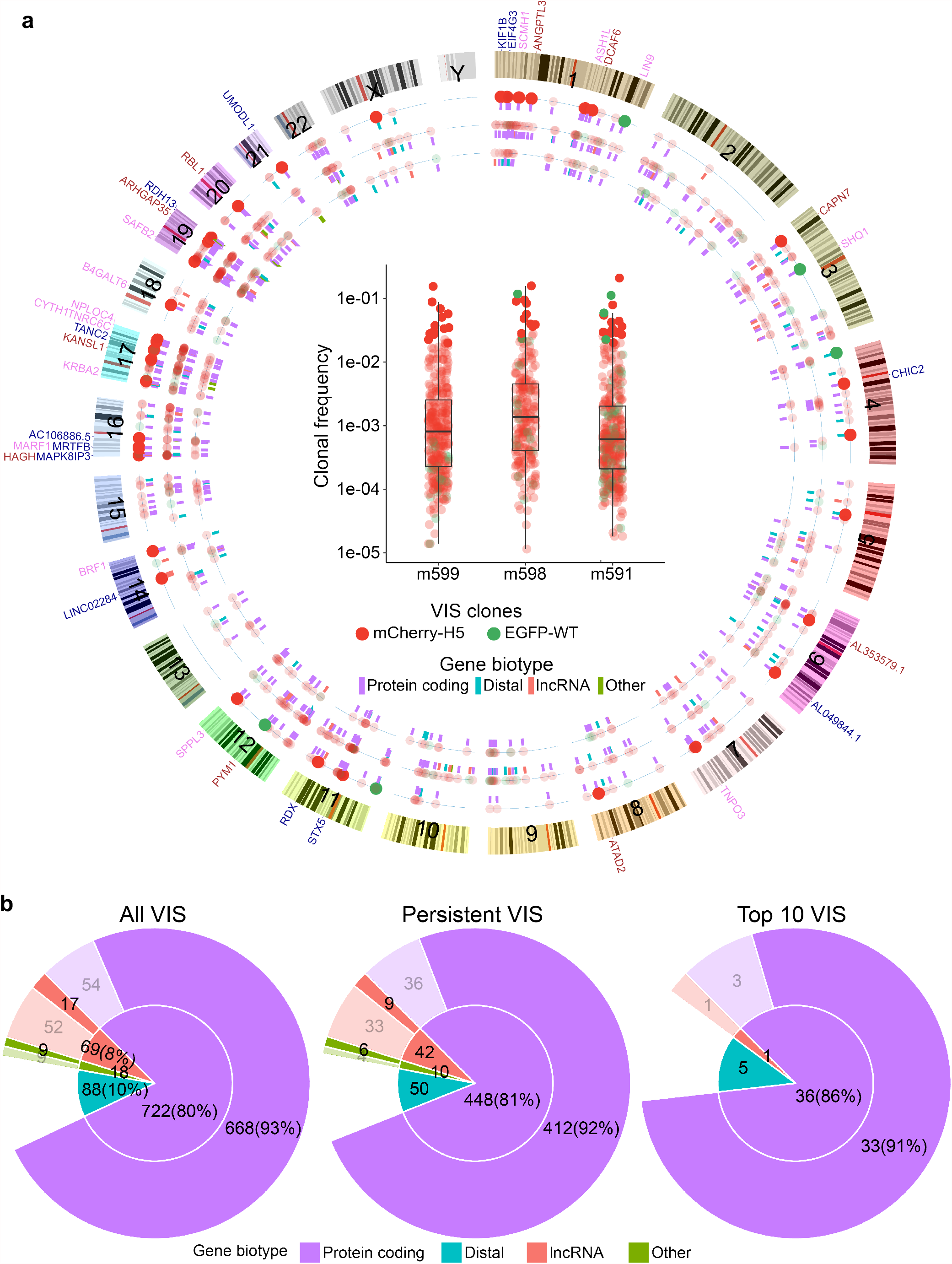
Chromosomal distribution of VIS and its bias for transcriptionally active genes: **a**) Circos plot shows genomic location of all 897 VIS from mice m599, m598, and m591. Box plots in the center show maximum frequency of mCherry-H5 (filled red dots) and EGFP-WT (filled green dots) VIS clones in mice m599, m598, and m591 over 6 weeks. Genomic location of mCherry-H5 (filled red dots) and EGFP-WT (filled green dots) VIS clones are plotted on three concentric circles depending on their maximum frequency over 6 weeks: Low frequency clones with maximum frequency below the 1^st^ quartile value (innermost concentric circle), High frequency clones above 3^rd^ quartile value (outermost concentric circle), and Medium frequency clones between 1^st^ and 3^rd^ quartile (middle concentric circle). Top 10 high frequency VIS clones are shown in darker colors. Color coded short line segments under each VIS show gene biotypes of VIS-proximal genes. Gene symbols above ideograms represent genes proximal to the top 10 VIS clones from mice m599 (blue), m598 (brown), and m591 (light pink). **b**) Classification of all, persistent, and Top 10 VIS clones based on biotype of VIS-proximal gene. Inner pie chart shows clones classified based on gene biotype of the most proximal gene. Outer Donut plots show number of VIS (in bracket % of VIS) proximal to active (dark color) or inactive (faded colors) genes. Active proximal genes have FPKM >1.

### Vector integration bias for transcriptionally active genes in human fetal liver HSPC

For in vivo repopulating clones we found VIS preference for genic regions in FL-HSPC however, the transcriptional state and expression level of the VIS-proximal genes prior to vector integration remains unknown. To address this, we analyzed transcriptomic (RNA-seq) and functional genomic (ATAC-seq and ChIP-seq) data from uncultured human FL-HSPC[30] isolated and processed using protocol identical to one used in our study (see Methods). Owing to the direct biological relevance of human FL-HSPC to our humanized BLT mice models, the multi-omics data is well suited to investigate the impact of vector integration on stemness and in vivo repopulation of vector-modified HSPC.

The gene expression (RNA-seq) data shows that of all in vivo detected clones, including the persistent clones and top 10 VIS clones, 694 out of 897 VIS (>77%) are within ±1Kb of transcriptionally active genes (FPKM>1) (Figure 4b outer donut chart). These VIS-proximal active genes include 668 protein coding genes, 17 lncRNA (non-coding genes) and 9 other genes (other non-coding genes and pseudogenes). This is significantly higher than the ∼27% of random IS proximal to active genes (p< 0.001, Supplementary figure 6A). The level of transcriptional activity of VIS-proximal genes (based on the FPKM value) was slightly higher than the median expression level of all active genes (Supplementary figure 5C). Moreover, the median gene activity level (FPKM) varied based on gene biotype, with protein coding proximal genes of all, persistent, and top 10 VIS clones having higher activity than proximal lncRNA or pseudogenes. Similar to in vitro observations using cell line and primary cells have reported vector integration bias for active genes with low to moderate expression [22, 46]. Our in vivo clonal tracking data show that for repopulating clones VIS bias for moderately active genes and against highly active genes. These findings suggest that similar to in vitro, in vivo viability and repopulation of gene-modified HSPC is likely linked to expression level of VIS proximal gene.

### Epigenetics of FL-HSPC reveals VIS bias for actively transcribed genes

We speculate that similar to expression activity of genes the chromatin structure of active genes strongly influences vector integration. Previous in vitro studies in cell lines suggested vector integration preference for genomic features such as select histone modification and DNase I hypersensitivity sites. However, neither such analysis is available for human FL-HSPC nor the influence of genomic location of VIS on in vivo engraftment of these HSPC has been explored. To investigate this, we analyzed functional genomic data of uncultured human FL-HSPC [30] with direct biological relevance to our humanized BLT mice models which are constructed by transplanting FL-HSPC in to NSG mice. To investigate whether vector preferred open chromatin or chromatin associated proteins, we analyzed chromatin accessibility (ATAC-seq) ChIP-seq data of 8 histone modifications H3K4me3, H3K9ac, H3K9cr, H3K27ac, H3K4me1, H3K79me2, H3K36me3, and H3K27me3 and RNApolII (chromatin associated RNA polymerase II), all from uncultured human FL-HSPC [30].

We explored the epigenetic landscape of all the VIS-proximal genes in FL-HSPC using the multi-omics data (Figure 5a profile plots and heatmaps). For active VIS-proximal genes (FPKM>1), our analysis showed active transcription markers such as open chromatin region (ATAC-seq peaks) and H3K4me3, H3K9ac, H3K9cr, and H3K27ac enriched regions near TSS as well as active enhancer markers (H3K27ac and H3K4me1 enriched regions). H3K36me3 and H3K79me2-enriched regions within the gene body of VIS-proximal active genes indicated genes being actively transcribed. We also detected the enrichment of RNApolII-bound chromatin (RNApolII) in active gene that coincided with TSS regions. For inactive (FPKM<1) VIS-proximal genes, these active gene marks were less prominent. Random IS-proximal genes showed similar enrichment profiles in active genes and their lack in inactive genes (Supplementary figure 6B). Overall, the epigenetic markers of the VIS-proximal genes show that the vector integration is biased for actively transcribed genes. However, further analysis was needed to determine presence of VIS bias for particular epigenetic features. We investigated possible enrichment of epigenetic features within ±1Kb region of VIS (Figure 5b). We found that vector integration did not occur near open chromatin (ATAC-seq) regions. Similarly, for histones H3K4me3, H3K9ac, H3K9cr, and H3K27ac high enrichment was not detected near VIS (Figure 5b lineplots), indicating that vector integration likely avoids transcription regulatory regions such as TSS and enhancer. The enrichment of RNApolII-bound chromatin (RNApolII) was also not visible near VIS further confirming that transcription regulatory regions are avoided. In contrast, enrichment of H3K36me3 and H3K79me2, near VIS, showing clear preference for actively transcribed regions within gene body. A Principal component analysis (PCA) on normalized enrichment levels (RPKM values) over a ±1Kb region of VIS (Figure 5b data) clearly separated H3K36me3 and H3K79me2 from all other features and showed no bias for random IS (Supplementary figure 7). These results further confirmed the VIS bias for actively transcribed regions.

**Figure 5:**
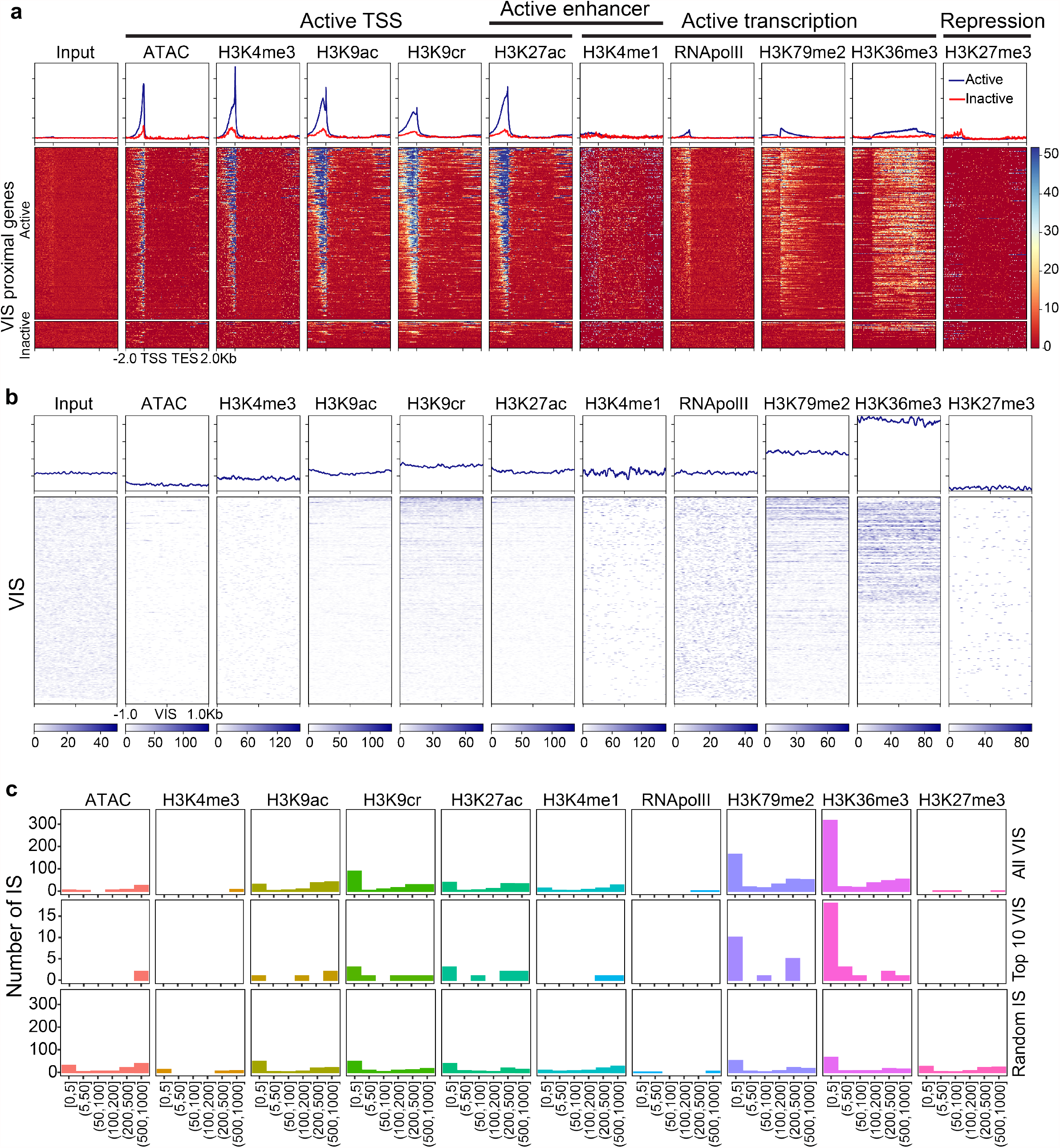
Epigenetic determinants of vector integration: **a**) Profile plots and heatmap for 10 chromatin features and input sample in active and inactive VIS-proximal genes in uncultured FL-HSPC. Profile plots show mean score for active (blue line) and inactive (red line) proximal genes. Score is calculated from normalized read count (RPKM) for each sample. Each row in heatmap shows expression level of 10 chromatin features in proximal genes from TSS to TES with 2Kb flanks upstream and downstream. Color scale key shows range of normalized expression. **b**) Profile plots and heatmaps for 10 chromatin features in region flanking ±1Kb of each VIS. Profile plots showing mean scores over ±1Kb region flanking VIS. Each row in the heatmap shows the expression level of 10 chromatin features in regions flanking ±1Kb of VIS. Individual color scale key shows the range of normalized expression for corresponding features. **c**) Bar plots show number of VIS within ±1Kb of enriched region (peak) of different chromatin features. VIS clones and random IS are binned by absolute distance in base pairs (bp) between the enriched region and IS. Bars show number of VIS in each bin. Top panel shows binning for all VIS, middle panel shows top 10 VIS clones, and bottom panel shows random IS falling within ±1Kb of enriched region.

### Vector integrations in repopulating clones were outside gene regulatory regions

In order to further investigate the proximity of VIS to epigenetic features at finer resolution, we called peaks for all the epigenetic markers (described in Methods) and measured the distance between the VIS and the center of the peak. Our analysis showed that 659 (73.5% of total) VIS were near (±1Kb) one of the 10 epigenetic features (Supplementary figure 8). Previous study using epigenetic data from different cell lines found a strong positive association of HIV integration to histones H3K4me1, H3K4me2, H3K4me3, and H3K9ac [47]. However, our analysis showed that only 6 out of 897 VIS were near H3K4me3, a marker for gene TSS, indicating disfavoring of transcription regulatory regions. Occurrence of only 63 VIS in H3K4me1 enriched regions indicating disfavoring of both H3K4 mono- and tri-methylation. Interestingly, compared to H3K4m1 and H4K4me3 more VIS associated with H3K9ac, H3K9cr, and H3K27ac (Supplementary figure 8A). Consistent with previous studies, we also observed that very few integration events (3 VIS) were near H3K27me3 enriched regions indicating that in repopulating clones vector integration happened outside of transcriptionally repressed regions (Figure 5b and 5c top panel, Supplementary figure 8). Analysis of open chromatin regions revealed that only 42 out of 897 VIS were in proximity of ATAC-seq peaks. Overall, fewer vector integration events happened in proximity of open chromatin (ATAC peak) or histones modification H3K4me3 and H3K4me1. These finding show that despite preference for active genes, in majority of repopulating clones, vector integration occurred outside the transcription regulatory regions (TSS and enhancers) of the genes. These finding also suggest integration in regulatory regions may adversely impacts survival of vector transduced FL-HSPC.

### Vector integrations predominantly occurred in H3K36me3 enriched regions

In repopulating clones, we observed bias against transcription regulatory regions of VIS-proximal genes despite significant VIS bias for active genes. While random IS were evenly distributed across all 10 histone marks (Figure 5c bottom panel, Supplementary figure 8C), in repopulating clones VIS distribution skewed towards H3K36me3 and/or H3K79me2 (Figure 5c top panel; Supplementary figure 8A-B). Out of 897 repopulating clones, 486 VIS (54% of total VIS) were within ±1Kb of histone marks H3K36me3, significantly higher compared to only 11.6% random IS being associated with the histone (p<0.001, Supplementary figure 6C). For the top 10 most frequent VIS clones, preference for all chromatin features (Figure 5c middle panel) was also similar to that for all VIS (Figure 5c top panel). A previous in vitro study using HIV based vector found that in human CD34+ cells, only 14.5% of vector integrations occurred within ±1Kb of H3K36me3 [23]. Compared to in vitro, we found 3.7 times more VIS to be within ±1Kb of H3K36me3 [23] for in vivo repopulating clones. Despite significant integration bias for active genes and in particular for histone H3K36me3, the clonal size distribution and repopulation of human cell in hu-BLT mice remained normal. These in vivo finding signify that both the vectors are safe and that vector integration near H3K36me3 appears less detrimental to engraftment and in vivo differentiation of vector transduced HSPC.

## Discussion

In this study we presented LoVIS-Seq, a new longitudinal clonal tracking method requiring a mere 25µl blood or less to monitor clonal behavior of gene-modified cells in small-animal models. LoVIS-Seq quantitatively captured the clonality of both control (mCherry-H1/H5) and anti-HIV (EGFP-WT) populations in whole blood. We provide the first-ever demonstration of simultaneous polyclonal repopulation of therapeutic vector-modified and control vector-modified cell populations in hu-BLT mouse blood. The polyclonal expansion resembles normal post-transplant HSPC clonal repopulation in mice [13], nonhuman primates [12, 17, 39] and humans [11, 40]. Notably, the clonal frequency data recapitulated the flow cytometric measurements. Persistent clones are major contributors in blood, BM, and spleen. The multi-omics data from uncultured FL-HSPC revealed that vector integration in VIS clones that repopulated in mouse environment is significantly biased toward H3K36me3 and/or H3K79me2 enriched regions. Remarkably, this vector integration bias appears inconsequential with respect to clonal repopulation, as gene-modified HSPC differentiated normally in vivo; this confirms the safety of our therapeutic and control lentiviral vectors.

We recently showed that in hu-BLT mice, monitoring clonal expansion of gene-modified and control populations within the same animal gives an unbiased analysis and allows a more direct assessment of therapeutic vectors [18]. In the current study, we longitudinally tracked both anti-HIV gene-modified and control vector-modified populations in the same hu-BLT mouse. Interestingly, we found a competitive growth pattern between two populations, with few clones from each population leading the expansion. The clonal profiles in both populations resemble typical after transplant clonal repopulation confirming safety of both vectors. Since safety and efficacy of multiple vectors can be tested in the same humanized mouse, LoVIS-Seq can reduce both cost and time of vector development.

Previous studies propose that in myeloablated mice, hematopoiesis tends to stabilize around 22 weeks post-transplant [15, 48] while a recent study suggested 16 to 24 weeks [49]; our data indicates clonal stabilization between 17 to 19 weeks post-transplant. Overall, the time scale of clonal repopulation of human HSPC in hu-BLT mice resembles that of mouse HSPC after transplant in mice. Cord blood HSPC transplanted in NGS mice also showed similar clonal behavior, with clonal stabilization starting near week 18 to 20 post-transplant [50, 51]. Although the timescales compare well with other studies, caution is due considering high incidence of graft versus host-disease related illnesses in xenograft mouse models.

HIV-1 and HIV-1 based vectors are known to favor transcriptionally active gene dense regions [44, 46], with preference for histone modification H3 acetylation, H4 acetylation and H3K4 methylation [47]. Some studies found no preference for DNase I hypersensitive sites [52] and H3K4 methylation being disfavored [53] and preference for H3K36me3 in Jurkat cells [53]. However, the implications of HIV-1 integration on cell fate are compounded by infection induced cytotoxicity. Another important study provides a link between HIV-1 integrations in genes related to viral replication and in vivo expansion and persistence of HIV-1 infected cells [45]. However, influence of epigenetic features proximal to HIV-1 IS on clonal expansion and persistence remained unexplored. The HIV-1 integration occurs with assistance from nuclear pore complex and targets the active genes closer to the nuclear pore and disfavor heterochromatin regions and active regions located centrally in the nuclease [54]. In partial agreement with previous studies, our analysis of repopulating VIS clones shows that clones with vector integration in transcriptionally active genes in FL-HSPC have higher probability of in vivo expansion however, RNA-Seq data indicates bias against highly expressed genes. For in vitro activated cord blood and BM derived CD34+ HSC, the lentiviral vector showed no preference for highly active genes [22]. This is likely due to either obstruction by transcriptional machinery or detrimental effect of vector integration on survival of the cell. Further investigation of 10 chromatin features showed that vector integration had strong bias toward actively transcribed regions marked by histone modifications H3K36me3 and H3K79me2. This bias could be attributed to LEDGF/p75, a chromatin binding protein essential for efficient HIV-1 integration [55, 56], that binds to integrases of HIV-1 [57, 58] and protects the pre-integration complex from degradation [59], whereas the N-terminal PWWP domain of LEDGF is known to preferentially interact with H3K36me3 [60, 61]. A recent study showed that inhibiting binding of LEDGF to integrases leads to increased distance between IS and H3K36me3 regions. H3K79me2 and H3K36me3 mark the gene body and H3K36me3 marks exons and is positioned near the 5’ end of the exon and is correlated with alternative splicing [62, 63]. Thus, VIS in proximity of H3K36me3 are likely to influence co-transcriptional splicing of the proximal gene as well as expression of the vector itself. Human HSPC clones reconstituting the mice blood have significant vector integration bias that suggest a link between in vivo differentiation of VIS clones and proximity of vector integration to H3K36me3. However, tracking of hundreds of single HSPC clones indicated that such transcription events have no adverse impact on the stemness of repopulating vector-modified HSPC.

Another recent study demonstrated use of CRISPR/Cas to introduce barcodes in the long-term HSPC and longitudinally tracked a very limited number of HSPC clones [51]. Comparatively, using LoVIS-Seq we have tracked ∼10 times more HSPC clones per animal with high accuracy and reproducibility. It is pertinent to note that to enable insertion of barcoded donor DNA into the host genome, HSPC need to undergo in vitro preconditioning and incubation before transplantation. The double stranded breaks introduced by CRISPR/Cas activate DNA damage responses causing significant delays in HSPC proliferation and affects their in vivo repopulation [64]. Additionally, off-target gene-editing by CRISPR/Cas remains a concern. In contrast, LoVIS-Seq does not require preconditioning of HSPC and provides a ready to use high-throughput clonal tracking assay for small-animal models. Furthermore, LoVIS-Seq has wider applicability owing to its adaptability to many lentiviral vectors commonly used to insert transgenes or reporter gene such as GFP.

LoVIS-Seq with whole genome amplification allows for quantitative assessment of clonal behavior in small-animal models. However, the accuracy and reproducibility of our assay depends on the initial number of cells used for MDA (Figure 1d). To minimize sampling errors, it is important to have a sufficient number of gene-marked human cells in each 25µl of blood or 10,000 cells to represent each clone in similar proportions as in the bulk population. Higher human reconstitution and gene marking are often desirable and necessary conditions wherein our assay provides optimal results.

## Conclusions

Our study revealed the in vivo dynamics of clonal expansion, emergence of stable stem cell clones, and consequences of vector integration bias on repopulation of human HSPC in mouse environment. Using a mere 25µl blood and LTR-indexed vectors, we provide first ever demonstration of dynamic polyclonal expansion in both control and therapeutic vector-modified populations in the same hu-BLT mice. Our study provides insights into clonal dynamics of human HSPC revealing a faster yet normal repopulation of human HSPC in mouse environment. In repopulating clones, we found significant VIS bias for actively transcribed regions specifically for H3K36me3-enriched regions indicative of vector integration in these regions being less detrimental to HSPC. These results provide a framework for understanding the clonal behavior of human HPSC repopulating in a mouse environment, critical for translating results from humanized mice models to the human settings. Moreover, our new assay provides an efficient tool for multifaceted analysis of clonal dynamics in murine and humanized-mouse models used extensively in infectious disease, cancer, gene-therapy, and stem cell research.

## List of abbreviations

HSPC: hematopoietic stem and progenitor cells
LoVIS-Seq: low volume vector integration site sequencing
BLT: Bone marrow-liver-thymus
FL: fetal liver
MDA: Multiple displacement amplification.

## Data availability

Raw RNA-seq, ATAC-seq, ChIP-seq data of uncultured FL-HSPC from published reference is available in Gene Expression Omnibus (GEO) with the accession code GSE111484.[30]

## Acknowledgements

We thank the technical support from the UCLA Center for AIDS research (CFAR) research cores including the CFAR Gene and Cellular Therapy Core and the CFAR Humanized Mouse Core. We would like also to thank Anna Sahakyan and Ruth Cortado for their help with making lentiviral vectors, transducing human CD34+ HSPC, and performing humanized mouse experiments.

