## Supplemental table and figures for "Longitudinal clonal tracking in humanized mice reveals sustained polyclonal repopulation of gene-modified human-HSPC despite vector integration bias"

| Sample | MDA | Amount of DNA used for VIS assay (µg) | Number of H1 clones | Number of WT clones |
|---|---|---|---|---|
| UT1 | No | 1 | 192 | 61 |
| UT2 | No | 1 | 221 | 94 |
| UT3 | No | 1 | 236 | 97 |
| 81000 Cells R1 | Yes | 1 | 234 | 102 |
| 81000 Cells R2 | Yes | 1 | 168 | 62 |
| 27000 Cells R1 | Yes | 1 | 229 | 95 |
| 27000 Cells R2 | Yes | 1 | 231 | 86 |
| 9000 Cells R1 | Yes | 1 | 214 | 78 |
| 9000 Cells R2 | Yes | 1 | 212 | 88 |
| 3000 Cells R1 | Yes | 1 | 209 | 74 |
| 3000 Cells R2 | Yes | 1 | 207 | 76 |
| 1000 Cells R1 | Yes | 1 | 199 | 89 |
| 1000 Cells R2 | Yes | 1 | 182 | 83 |

Supplementary table 1: Amount of unamplified DNA and MDA amplified DNA used for VIS .

| m599 | | | | m598 | | | | m591 | | | |
|---|---|---|---|---|---|---|---|---|---|---|---|
| **Week** | **hCD45 %** | **EGFP %** | **mCherry %** | **Week** | **hCD45 %** | **EGFP %** | **mCherry %** | **Week** | **hCD45 %** | **EGFP %** | **mCherry %** |
| 13 | 40 | 0.35 | 88.4 | 13 | 28.5 | 2.54 | 58.7 | 13 | 24.7 | 10.8 | 81 |
| 15 | 67.2 | 0.19 | 88.8 | 15 | 33.7 | 5.33 | 56.1 | 15 | 42.4 | 25.7 | 65.8 |
| 17 | 46.1 | 0.12 | 74.9 | 17 | 58.6 | 10.8 | 34.1 | 17 | 69.6 | 39.5 | 43.2 |
| 19 | 57.6 | 0.12 | 72.1 | 19 | 73.9 | 3.05 | 49.7 | 19 | 60.9 | 32.1 | 55 |

Supplementary table 2: Flow cytometry data from three animals (m599, m598, and m591) for week 13, 15, 17 and 19. Table shows percentage of EGFP+ or mCherry+ cells within human CD45 cells (hCD45+) in blood.

| Sample | MDA | Amount of DNA used for VIS assay (ug) | | |
|---|---|---|---|---|
| | | m599 | m598 | m591 |
| Whole blood ~1ml (Week 19) | No | 2 | 1.7 | - |
| Bone marrow (Week 19) | No | 2 | 2 | 2 |
| Spleen (Week 19) | No | 2 | 2 | 2 |
| Week 13 R1 (25 ul PB) | Yes | 2 | 2 | 2 |
| Week 13 R2 (25 ul PB) | Yes | 2 | 2 | 2 |
| Week 15 R1 (25 ul PB) | Yes | 2 | 2 | 2 |
| Week 15 R2 (25 ul PB) | Yes | 2 | 2 | 2 |
| Week 17 R1 (25 ul PB) | Yes | 2 | 2 | 2 |
| Week 17 R2 (25 ul PB) | Yes | 2 | 2 | 2 |
| Week 19 R1 (25 ul PB) | Yes | 2 | 2 | 2 |
| Week 19 R2 (25 ul PB) | Yes | 2 | 2 | 2 |

Supplementary table 3: Amount of unamplified DNA and MDA amplified DNA used for clonal tracking assay for mouse m599, m598 and m591.

| m599 | | | |
|---|---|---|---|
| Week | Human Cells/µl blood | Number of cells/MDA replicate | # of VIS clones |
| 13 | 742 | 18558 | 317 |
| 15 | 1018 | 25461 | 280 |
| 17 | 1329 | 33230 | 274 |
| 19 | 403 | 10070 | 276 |
| m598 | | | |
| Week | Human Cells/µl blood | Number of cells/MDA replicate | # of VIS clones |
| 13 | 960 | 24002 | 200 |
| 15 | 726 | 18156 | 193 |
| 17 | 1125 | 28118 | 188 |
| 19 | 625 | 15620 | 182 |
| m591 | | | |
| Week | Human Cells/µl blood | Number of cells/MDA replicate | # of VIS clones |
| 13 | 533 | 13336 | 277 |
| 15 | 819 | 20474 | 250 |
| 17 | 1271 | 31766 | 260 |
| 19 | 619 | 15480 | 226 |

Supplementary table 4: Number of blood cells used for MDA amplification and number of VIS clones detected at each time ploint.

Supplementary figure 1: A) Experimental protocol to estimate minimum number of cells required for MDA to reproduce VIS clonal profile from unamplified DNA of bulk BM cells. (B) Scatter plot showing the correlation between replicate R1 and R2 of unamplified samples (i), between unamplified replicate R1 and amplified DNA from 81000 cells (ii) and between unamplified replicate R1 and amplified DNA from 1000 cells (iii). Each dot in scatter plot is a VIS clone (red dot for mCherry-H1 clones and green dots show EGFP-WT clones). Correlation coefficient values in red show Pearson's r for mCherry-H1 clones and in green for EGFP-WT clones. C) Stacked bar plots of Clonal profiles for all the clones (mCherry-H1 and EGFP-WT) D) Top panel shows clonal profiles for mCherry-H1 VIS and clonal profiles of EGFP-WT VIS are shown in the bottom panel. VIS clones are arranged in descending order (bottom to top) of their frequency in unamplified replicate R2. Same order of VIS clones is maintained for all the samples. E-F) Plot showing Pearson correlation between all the unamplified and MDA amplified DNA samples. Size of circle is proportional to Pearson's r value. Color of the circle corresponds to Pearson's r value as given in color scale key and digits in yellow are Pearson's r values. G) Dot plot showing number of mCherry-H1 VIS (red) and EGFP-WT VIS (green) detected in different unamplified replicates and MDA amplified replicates for different number of cells. H) Barplot showing ICC values for 3 replicates of unamplified DNA and between 2 MDA amplified DNA replicates and 3 replicates of unamplified DNA for different number of cells used for MDA.

I) Heatmap plot showing average correlation (top Pearson) between MDA amplified DNA replicates and unamplified DNA replicates calculated using different number of top high frequency clones. The bottom heatmap showing the percentage (100 x color key value) contribution by the different number of top n high frequency clones to the gene modified cell population.

Supplementary figure 2: A) Flow cytometry data of mCherry and EGFP vector transduced and mock transduced FL-CD34 cells. Details of the experiments are provided in methods section in the main manuscript. B) Line plots showing hCD45+, B, and T cell count at different weeks post-transplant in mice m599, m598, and m591.

Supplementary figure 3: High reproducibility between MDA replicates. A) Each scatter plot showing VIS frequencies detected in two replicates at each timepoint of m599, m598, and m591. B) Scatter plot showing correlation between clonal frequency by VIS assay and corresponding % gene marking by flow cytometry.

Supplementary figure 4: Clonal evolution over time. A) Heatmaps showing percentage decrease in shared clone between two timepoints in mCherry-H5 (top plot) and EGFP-WT (bottom plots) population. Digits inside tiles on the diagonal with white background show number of VIS detected at each time point. Colors of each heatmap tile corresponds to percentage of clones shared and color key is provided on the right. Digits in each tile show number of VIS shared between two timepoint. B) Heatmaps showing change in correlation of clonal profiles overtime. Color and size of the squares in represents Pearson's r values (shown in yellow digits). Color of square corresponds to the color key on the righthand side. C) Barplots showing Shannon and Simpson index values at different weeks post-transplant.

Supplementary figure 5:  Chromosomal distribution of VIS and activity status of proximal genes. A) Barplot showing number of VIS in each chromosome and number on top of each bar shows percentage of VIS present in that chromosome. B) Gene ontology analysis of persistent clone proximal human genes and mouse orthologs. Results showing significantly (p value <0.01) enriched biological process predicated using proximal human genes by DAVID Gene Functional Classification Tool web-based tool. c) Boxplot showing FPKM values estimated from RNA-seq of FL-HSPC cells for three replicates. FPKM values are plotted for all genes, all VIS proximal genes, persistent VIS proximal genes, and top 10 VIS proximal genes. Gene are classified and color coded by their ensembl gene biotype defined in legends. Other genes exclude lncRNA, Protein coding, and pseudogenes.

Supplementary figure 6: Distribution of Random IS based on proximity to gene. A) Inner pie chart show classification of based on gene classification of its proximal gene. Outer Donut plots shows number of (% of) VIS with active (dark color) inactive (faded colors) proximal genes. Active proximal genes have FPKM >1.  B) Profile plots and heatmap for 10 chromatin features and input sample in active and inactive random IS proximal genes in uncultured FL-HSPC. Profile plots show mean score of active (blue line) and inactive (green line) proximal genes. Score is calculated from normalized read count (RPKM) for each sample. Each row in heatmap show expression level of 10 chromatin features in proximal gene from TSS to TES with 2Kb flanking upstream and downstream. Color scale key shows range of normalized expression. C) Profile plot and heatmap for 10 chromatin features in region flanking ±1Kb of each random IS. Profile plots showing mean scores over region flanking ±1Kb of random IS. Each row in heatmap show expression level of 10 chromatin features in region flanking ±1Kb of random IS. Individual color scale key shows range normalized expression for corresponding feature.

Supplementary figure 7: Principal component analysis on normalized read count over region flanking ±1Kb of VIS (A) and random IS (B). Top PCA plots showing eigen values for PC1 and PC3 and in bottom Scree plot bar showing % variance explained by each PC and line plot cumulative variance explained. PCA analysis was done using software deeptools.

Supplementary figure 8: Distribution of VIS across epigenetic landscape in fetal liver CD34+ cells. A) Tile plot showing number of VIS proximal (within ±1kb) to an epigenetic feature (tiles on the diagonal). Values in tiles above diagonal show number of VIS that were fell with ±1Kb of other epigenetic features. B) Heatmap showing distance (in bp) of all 897 VIS (each row) from nearest gene and 10 epigenetic features. VIS are arranged bottom to top based on activity status of gene (active FPKM>1 or inactive genes) and distance from gene. C) Tile plot showing number of random IS proximal (within ±1kb) to an epigenetic feature (Tiles on the diagonal). Values in tiles above diagonal show number of random IS that were fell with ±1Kb of other epigenetic features.

Supplementary figure 9: Detailed LoVIS-seq workflow. A) Protocol for isolation cells from 100µl mouse blood. Transfer blood in microcentrifuge tubes (1.5 ml) and spin tubes at 2400rpm for 5 min. Remove plasma resuspend cells in 50µl of flow cytometry anti-body master mix in staining buffer and incubated for 30min at RT. After incubation add 1ml red blood cell lysis buffer and incubate for 5 min at RT. Spin the cells remove supernatant and resuspend cells in 1ml sainting buffer. Spin the cell and remove supernatant. Resuspend the cells in 16µl of 1XPBS provided with REPLI-g Single Cell Kit from Qiagen (Cat #150343). Out of 16µl, use 8µl to make two replicates by transferring it into 2 tubes (equivalent to 25µl blood). Store tubes at -20C or can be used for MDA amplification. Rest of 8µl is resuspended in 300µl of fixed buffer and use for flow cytometry. B) Protocol for performing MDA directly on cells. Thaw cells if stored at -20C. Add 3µl of denaturation buffer (Protocol preparing denaturation buffer is provided with REPLI-g Single Cell Kit) to each 4µl replicates. Incubate at 65C for 10 min then add 3µl of stop solution (provided in REPLI-g Single Cell Kit) to each tube. Add 40µl of reaction mix (Protocol preparing reaction mix is provided with REPLI-g Single Cell Kit) to each tube and incubate for 8hr at 30C. After 8hr, to stop the reaction incubate tubes at 65C for 3min. Tubes can be stored in -20C or can be used for DNA purification. C)Protocol for purification of amplified DNA. Add 150µl of 1XPBS to the tube containing 50µl of amplified DNA (final product from previous step). Add 200µl buffer AL provided with QIAamp® DNA Mini Kit (Cat# 51304) and vortex for 15s and spin briefly. Add 200µl of ethanol (>96%) and vortex for 15s and spin briefly. Transfer the content to QIAamp spin column. Centrifuge at 8000rpm for 1min, discard the flow through. Add 500µl of buffer AW1 (provided with QIAamp® DNA Mini Kit) to columns. Centrifuge at 8000rpm for 1min, discard the flowthrough. Add 500µl of buffer AW2 (provided with QIAamp® DNA Mini

Kit) to columns. Centrifuge at 14000rpm for 2min, discard the flowthrough.  Transfer columns to new microcentrifuge tubes (1.5 ml). Add 200µl of buffer AE (provided with QIAamp® DNA Mini Kit) to columns. Centrifuge at 8000rpm for 1min. Flowthrough in the tubes contains purified amplified DNA that can be stored -20C for long term. The purified DNA is used for VIS assay described in detail at the end of the supplementary text.

Supplementary figure 1

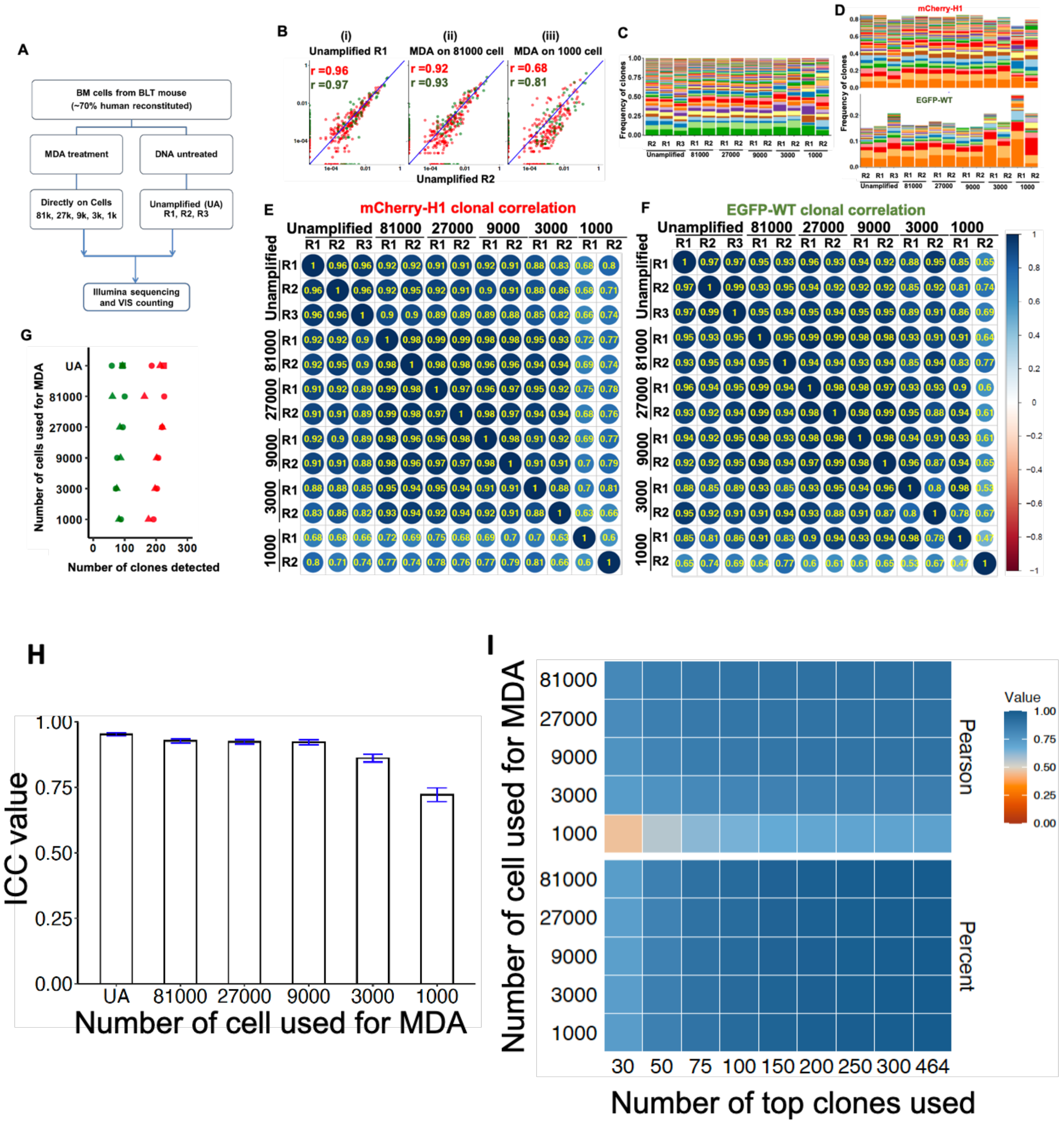

Supplementary figure 2

A

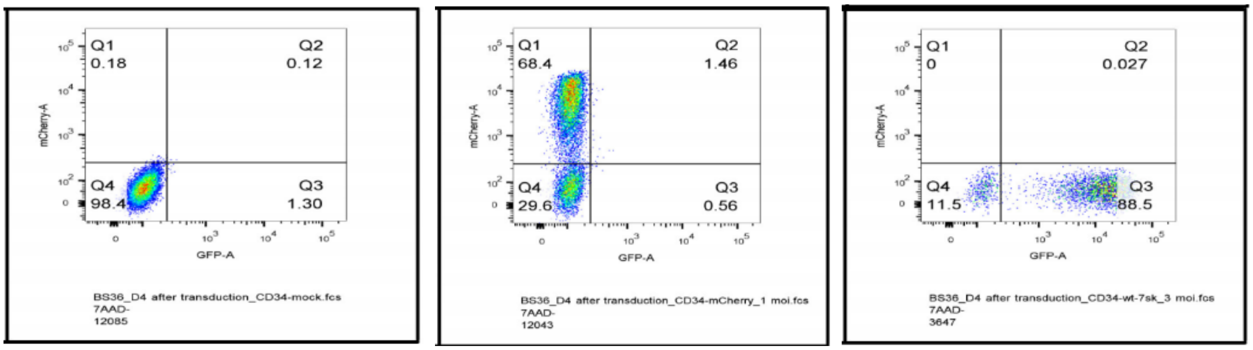

Mock transduction

mCherry transduction MOI=1

EGFP transduction MOI=3

B

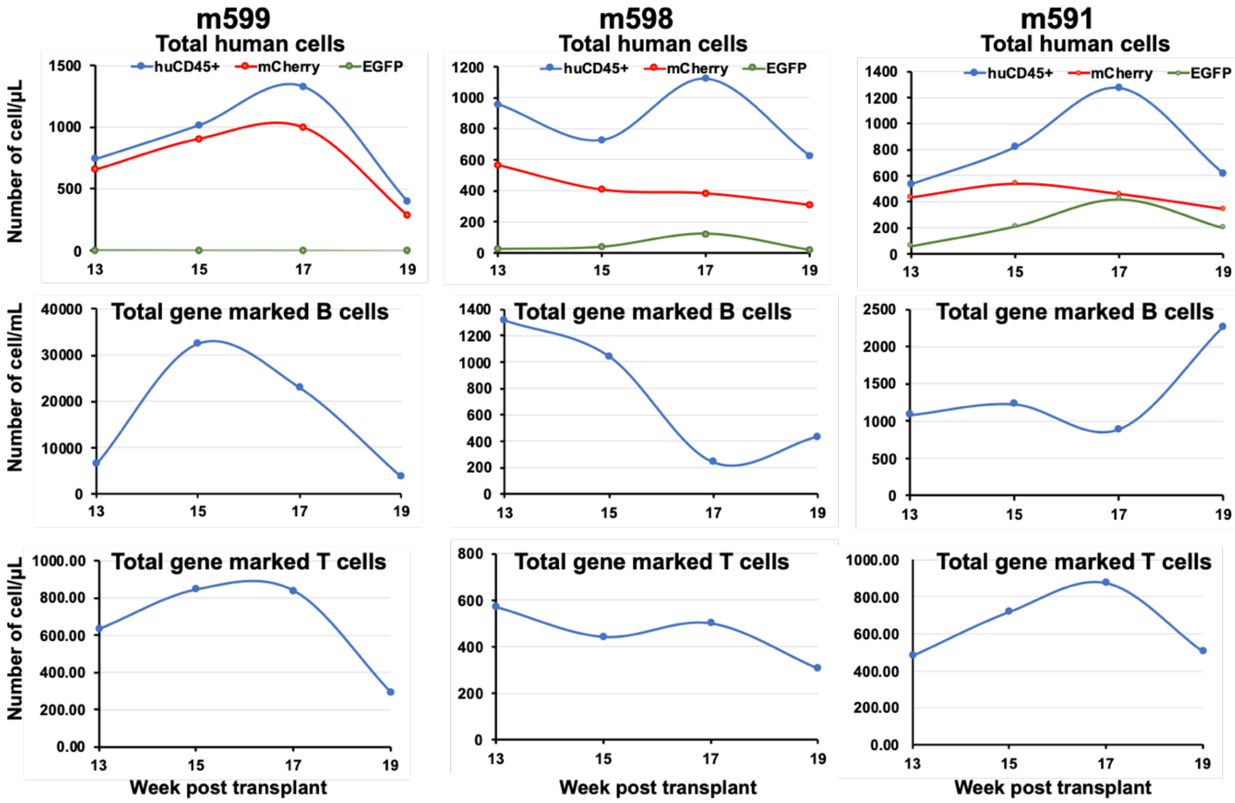

Supplementary figure 3

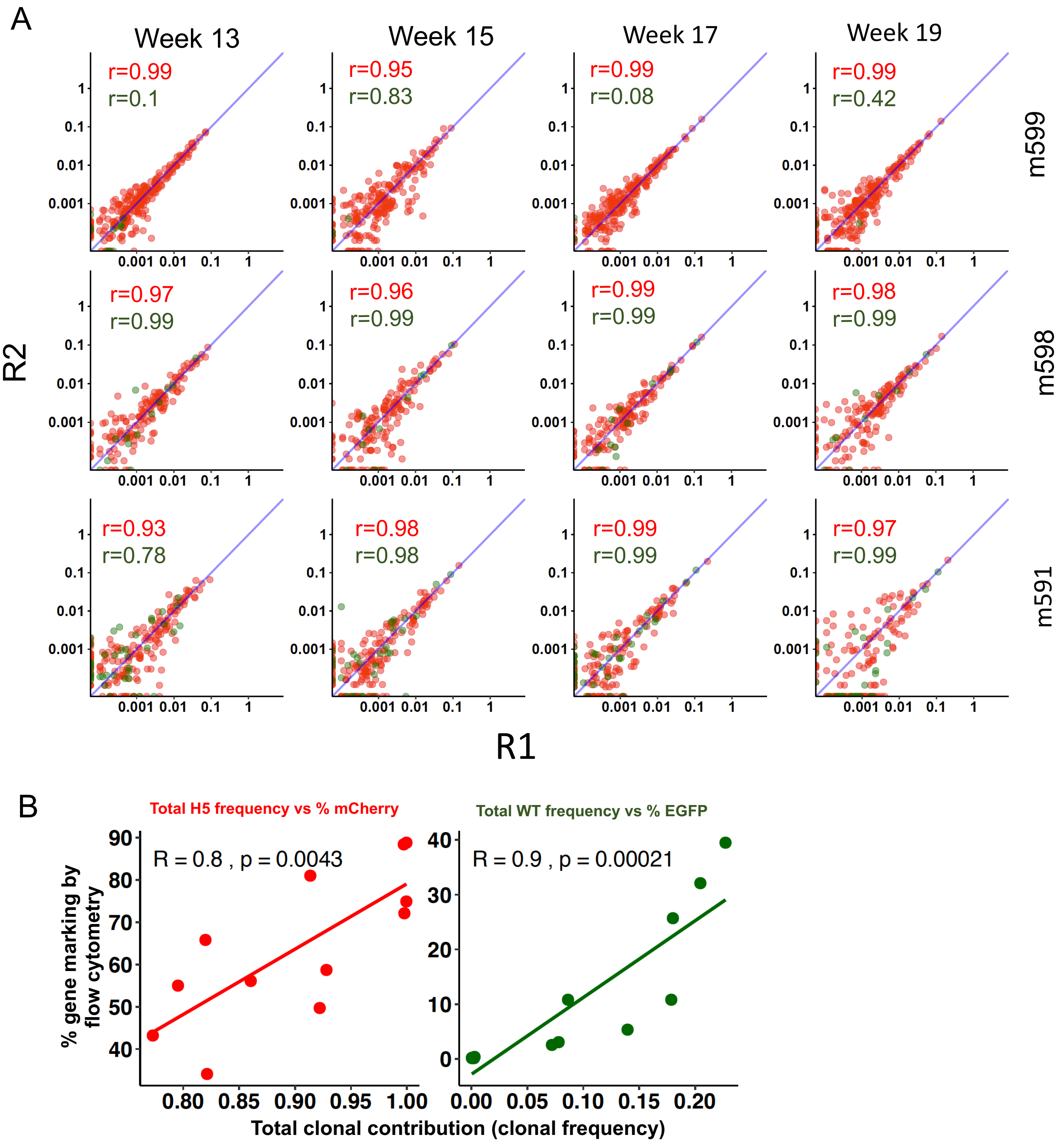

Supplementary figure 4

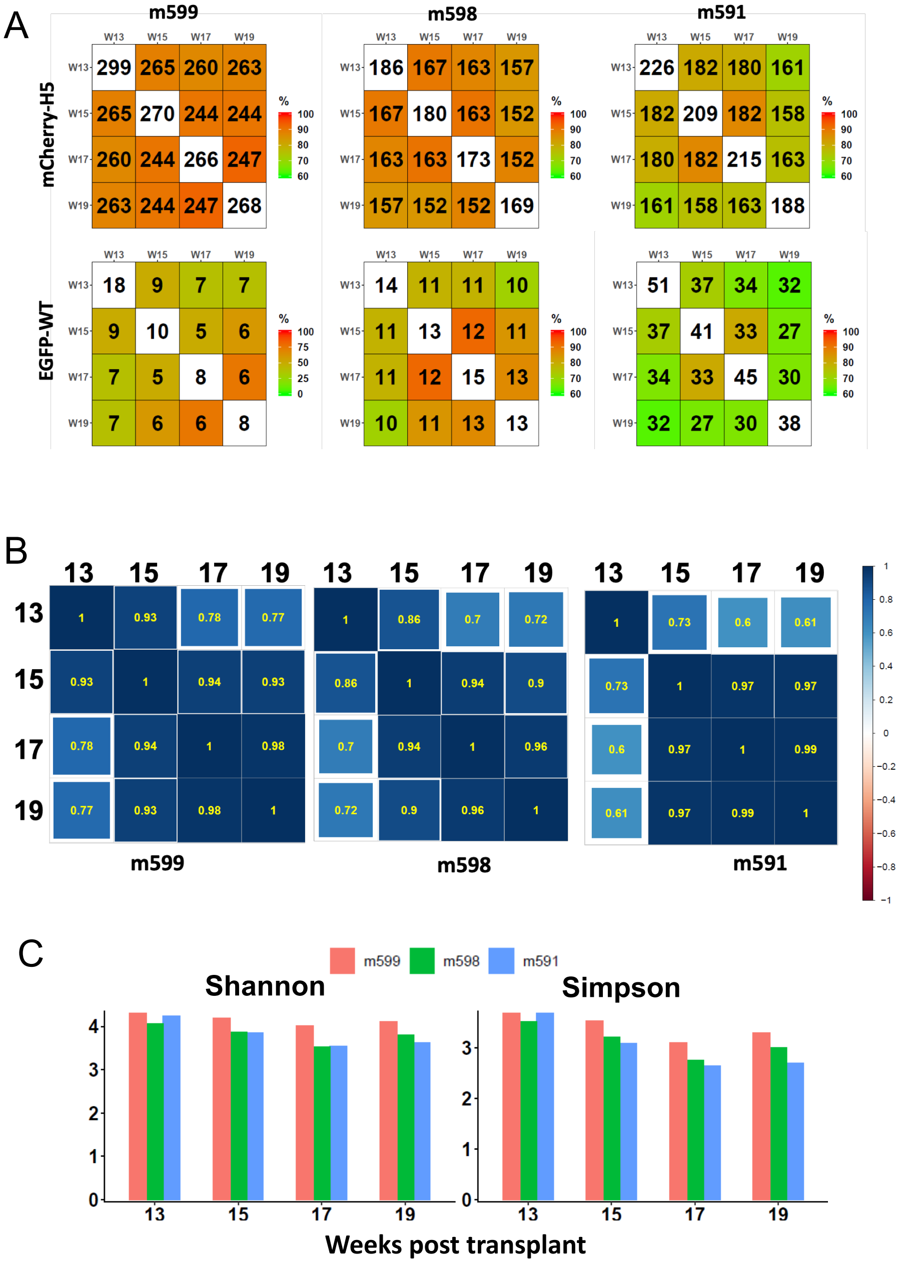

Supplementary figure 5

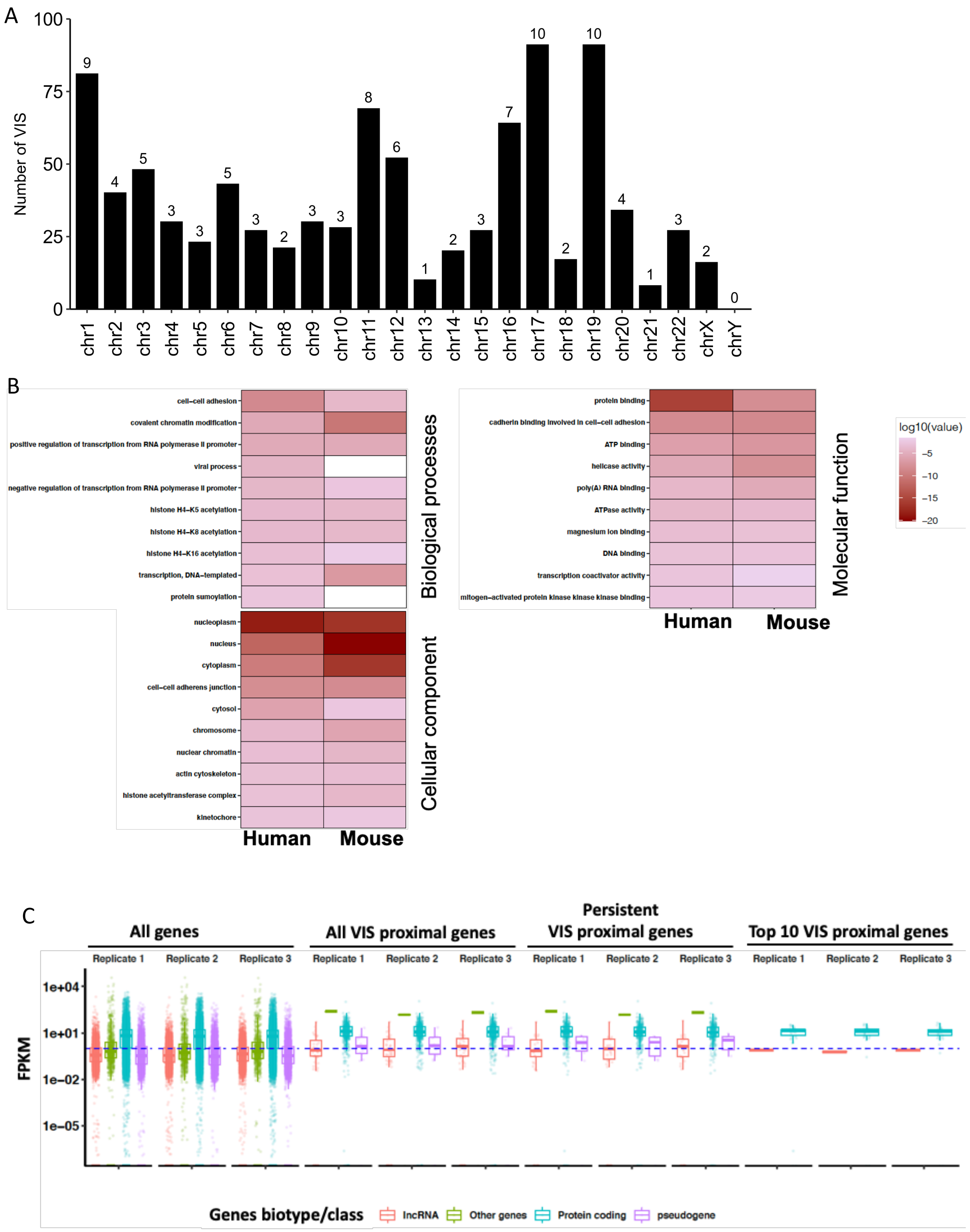

Supplementary figure 6

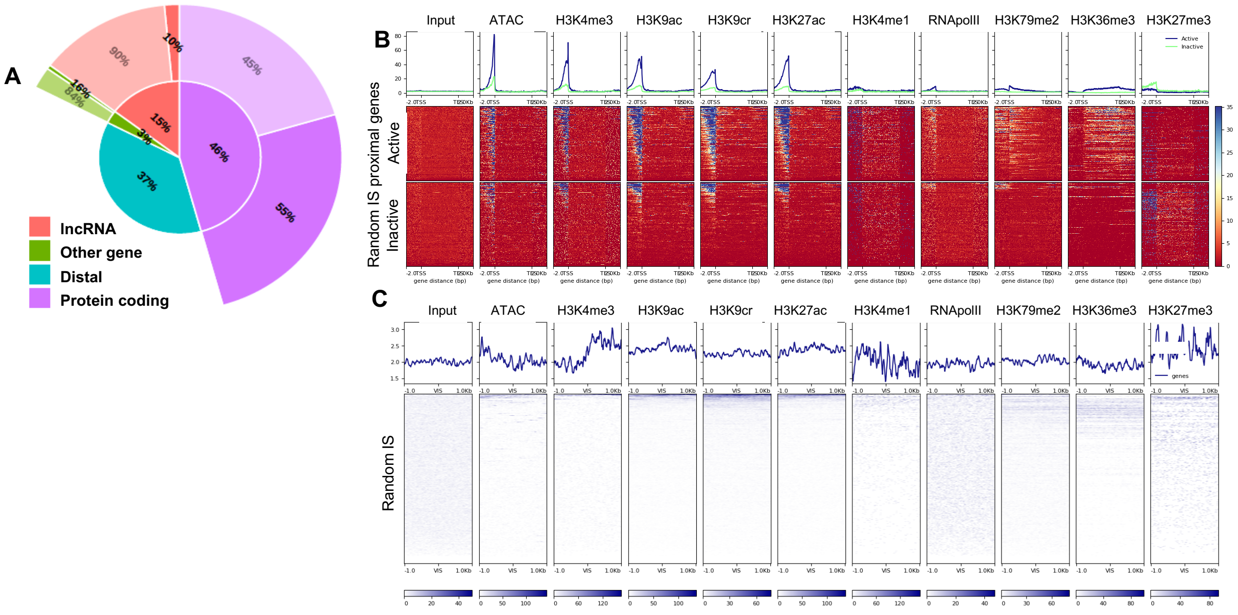

Supplementary figure 7

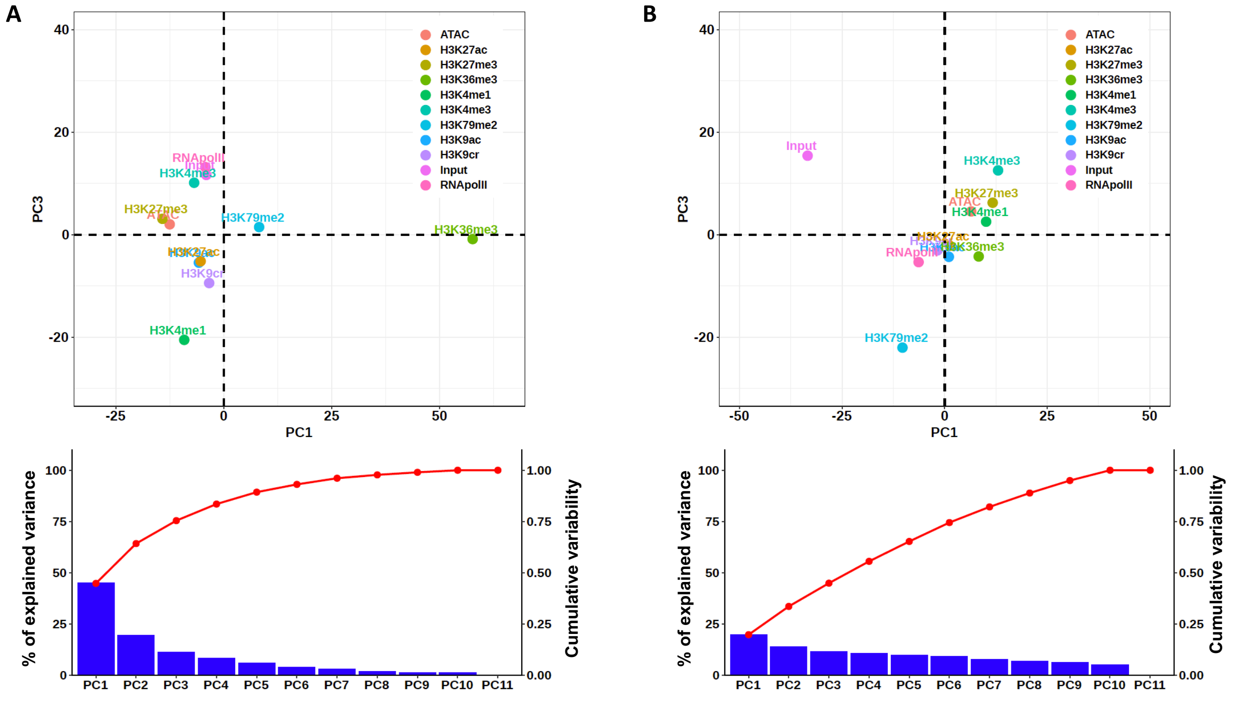

Supplementary figure 8

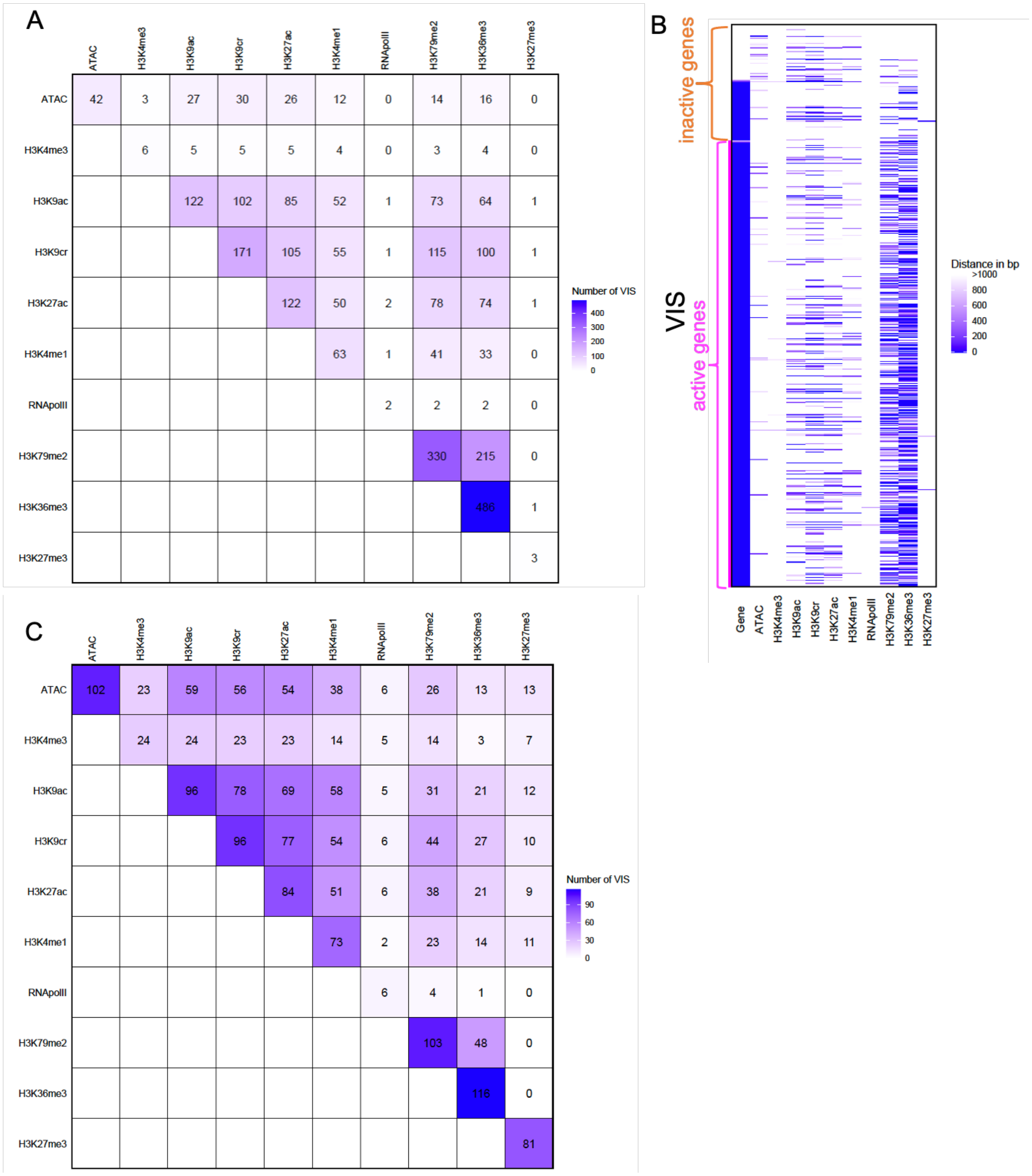

Supplementary figure 9

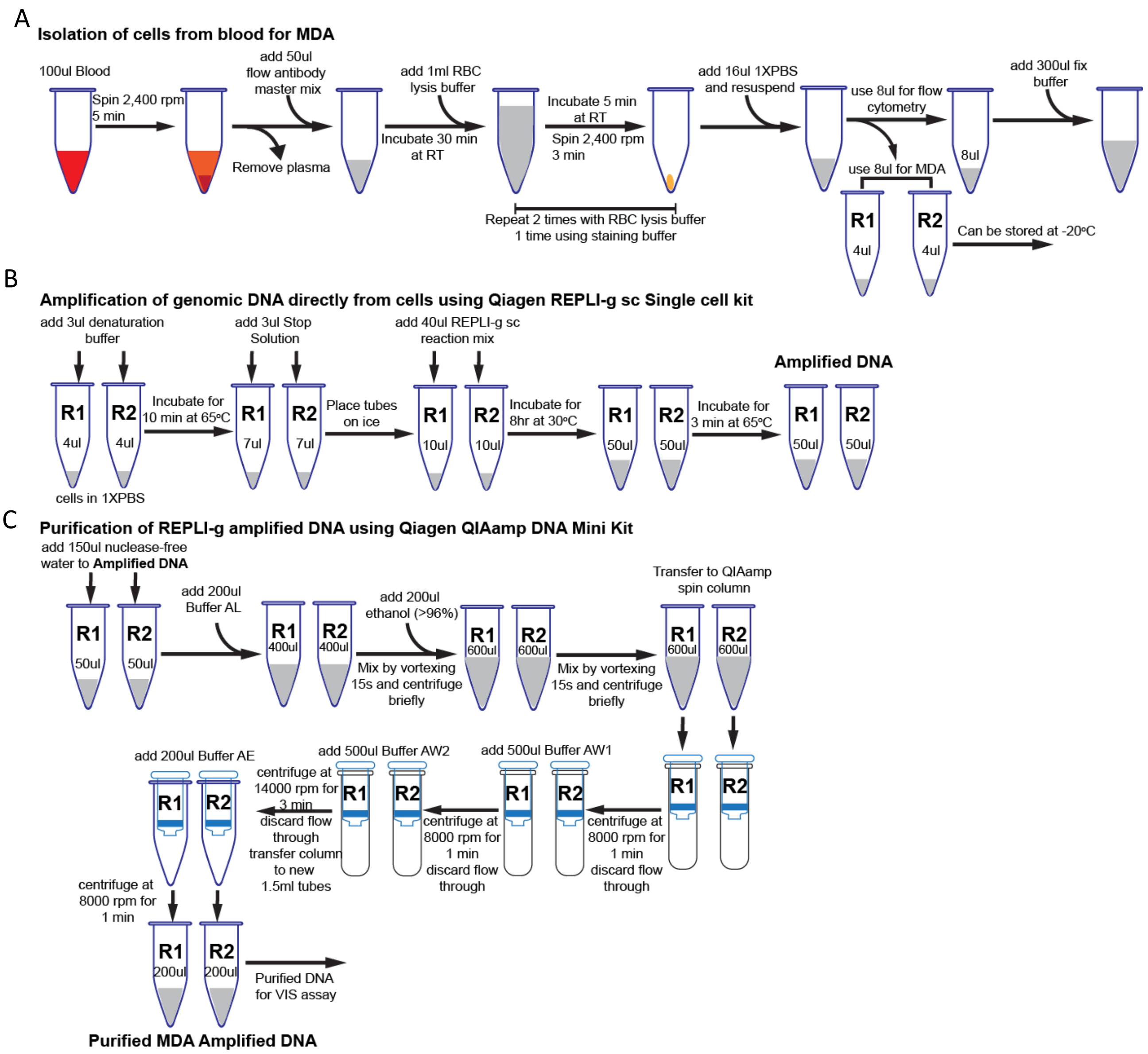

Detailed protocol of VIS assay:

### Genomic DNA denaturation & primer extension

Total reaction in 200ul

| | |
|---|---|
| Genomic DNA or Purified MDA amplified DNA | 1-2ug |
| Primer1 100mM | 0.25 |
| Primer2 100mM | 0.25 |
| 10X picomaxx buffer (invitrogen) | 20 |
| 10mM of deoxynucleoside triphosphates | 4 |
| picomaxx enzy | 2.5 |
| H2O | ? |
| total | 200ul |

/5BiotinTEG/CA GAT CTG AGC CTG GGA GCT C

/5BiotinTEG/CT GGC TAA CTA GGG AAC CCA CT

Aliquot into 66ul into each PCR tube

**Condition for extension**

94°C for 5 minutes

56°C for 3 minute

72°C for 5 minutes

4C

**DNA purification**

| |
|---|
| Pull 3 tubes DNA together |
| Qiagen PCR purification |
| Elute with 50ul 1:5 EB |

### RsaI digestion

| | micro-litters |
|---|---|
| purified DNA | 50ul |
| 10X NEB CutSmart buffer | 10 |
| RsaI (NEB, activity may differ by Lot#) | 2 |
| H2O | 38 |
| total | 100ul |

incubate at 37 C for 1 hour

| **CviQI digestion :** | micro-litters |
|---|---|
| CviQI (NEB, activity may differ by Lot#) | 1ul |
| add into the step 3 reaction | |
| incubate at 25 C for 45min | |

| **Blunt ending : 1** | micro-litters |
|---|---|
| 10mM dNTPs | 0.4 |
| DNA Polymerase I large (klenow) fragment | 0.5 |
| add into the step 4 reaction | 0.9ul |
| incubate at 25 C for 1 hour | |

**Streptavidin bead binding : 3~4 hours**

| |
|---|
| Prepare beads as follows, for each sample |
| Take 50ul beads /sample, |
| Remove the supernatant using the magnetic stand |
| Add 200 ul of TE buffer with 1M NaCl and voltex shortly |
| Spin at 1,500 rpm for 1 min |
| Remove the supernatant using the magnetic stand |
| Add 100ul "Binding Buffer" to the bead |
| Mix by pepitting (No votex) |

Add 100ul of prepared beads to the tube from step 5 (101.9ul)

Incubate at room temperature for 2~3 hours (vortex frequently or incubate samples on the rotating wheel)

**DNA linker preparation**

While step 6, prepare DNA linkers

| **Linker master Mix ( use 0.5ul/sample below)** | micro-litters | |
|---|---|---|
| 100uM of BL-L-Link-A oligos* = BHLinkA(HPLC purified) | 1 | CGGATCCCGCATCATATCTCCAGGTGTGACAC |
| 100uM of Blunt Link-s oligos* = BluntLinkS | 1 | CAC CTG GAG ATA TGA TGC GGG ATC CG |
| 5M NaCl | 1 | |
| H2O | 2 | |
| total | 5 | |

Incubate at 95 C for 5 mins and cool down slowly.

* Oligos for blunt-ended linkers - see below for the sequences

### Linker ligation : 3 hours ~ overnight

After the step 6, wash beads as follows
Spin at 1,500 rpm for 1 min
Remove the supernatant using the magnetic stand

Add 200 ul of "washing buffer" from the kit and vortex shortly
Spin at 1,500 rpm for 1 min
Remove the supernatant using the magnetic stand

Add 200 ul of 1:5 AE
Spin at 1,500 rpm for 1 min
Remove the supernatant using the magnetic stand

Add 200 ul of "1X NEB T4 DNA ligase buffer" and vortex shortly
Spin at 1,500 rpm for 1 min
Remove the supernatant using the magnetic stand

add the "ligation reaction solution-see below-" to the beads

| **Ligation reaction solution:** | **ul** |
|---|---|
| 10X NEB T4 DNA ligase buffer | 10 |
| 5X Invitrogen T4 DNA ligase buffer | 20 |
| $H_2O$ | 165 |
| linker solution (from the step 7) | 0.5 |
| NEB T4 DNA ligase | 5 |
| total | 200 |

Add 200ul of Ligation solution to beads

Put the tube on the rotating wheel and incubate it at room temparature (22 C) for 3 hours or overnight

### Wash beads

wash beads as follows
Spin at 1,500 rpm for 1 min
Remove the supernatant using the magnetic stand

Add 500 ul of 1:5 AE and vortex shortly

Spin at 1,500 rpm for 1 min
Remove the supernatant using the magnetic stand

Add 800 ul of "1X Taq Pol PCR buffer" and vortex shortly
Spin at 1,500 rpm for 1 min
Remove the supernatant using the magnetic stand

Add 50ul of "1X Taq Pol PCR buffer" and ready for 1st PCR test

**The beads can be stored at 4C for a few days**

**The 1st PCR**

use all remaining DNA beads

To amplify both the left and right vector- cellular DNA junctons

| | micro-litters | |
|---|---|---|
| DNA beads | 40 | |
| 100uM of primer1 | 2 | CTG GCT AAC TAG GGA ACC CAC T |
| 100uM of linker primer L1 | 2 | GTGTCACACCTGGAGATAT |
| 10mM DNTP | 4 | |
| picomaxx buffer (invitrogen) | 20 | |
| picomaxx | 8 | |
| H2O | 124 | |
| | 200 | |

aliquote into 66ul into each PCR tube

PCR condition

| | | |
|---|---|---|
| 94°C | 2 min | |
| 94°C | 20 sec | |
| 56°C | 25 sec | 25 cycles |
| 72°C | 2 min | |
| 72°C | 5min | |
| 4°C | | |

**Qiagen PCR Clean-up kit**

Qiagen PCR cleanup procedure

Elute 50ul 1:5 EB, (possible another 50ul EB elution, save in case)

**You can use restriction enzymes to remove "internal control" bands-using SfoI**

Digest 1st PCR product with SfoI before the 2nd PCR to remove "internal vector"

100ng/20ul for SfoI digestion

| | | |
|---|---|---|
| DNA | 10ul | *=100ng 1st PCR PRODUCT* |
| NEB CutSmart buffer | 2ul | |
| SfoI | 1ul | |
| H2O | 7ul | |
| | 20ul | 37C 1hour |

will use 5ul for second Right PCR

**2nd PCR right vector-cellular DNA junction**

| | | |
|---|---|---|
| 1st PCR product | 5ul | *=after sfoi digestion* |
| 100uM primer | 0.5ul | ACT CTG GTA ACT AGA GAT CC |
| 100uM primer L2 | 0.5ul | GGA GAT ATG ATG CGG GAT C |
| 10mM DNTP | 1ul | |
| picomaxx enzyme (invitrogen) | 2ul | |
| 10X picomaxx buffer | 5ul | |
| H2O | 36ul | |
| total | 50ul | |

PCR condition

| | | |
|---|---|---|
| 94°C | 1 min | |
| 94°C | 20 sec | |
| 58°C | 25 sec | 22 |
| 72°C | 2 min | |
| 72°C | 5min | |
| 4°C | | |

Run 5ul of PCR product on Qiaexel

** Test cycle numbers for 2nd PCR

*** Test sfoI digestion after determining the cycle number

**<u>Qiagen PCR Clean-up kit for Right</u>**

Qiagen PCR purification

Elute 50ul of 1:5 EB

Product of Second PCR is used for sequencing after attaching Illumina seqeuncing primers by PCR
